# Neonatal levels of acute phase proteins and risk of autism spectrum disorders

**DOI:** 10.1101/2020.02.13.947572

**Authors:** Renee M. Gardner, Brian K. Lee, Martin Brynge, Hugo Sjöqvist, Christina Dalman, Håkan Karlsson

**Affiliations:** Department of Global Public Health, Karolinska Institutet, Stockholm, Sweden, 17177; Department of Epidemiology and Biostatistics, Drexel University School of Public Health, Philadelphia, PA, USA; A.J. Drexel Autism Institute, Philadelphia, PA, USA; Centre for Epidemiology and Community Medicine, Stockholm County Council, Stockholm, Sweden, 17129; Department of Neuroscience, Karolinska Institutet, Stockholm, Sweden, 17177

**Author notes:** Indicates that the authors made an equal contribution to the work (Christina Dalman, Håkan Karlsson). **Corresponding Author**: Renee Gardner, Department of Global Public Health (K9), Karolinska Institutet, SE-171 77 Stockholm, Sweden.

**Keywords:** Autism spectrum disorders, acute phase protein, innate immunity, inflammation, infection, C reactive protein, ferritin

## Abstract

**Background:** Immune signaling pathways influence neurodevelopment and are hypothesized to contribute to the etiology of autism spectrum disorders (ASD). We aimed to assess risk of ASD in relation to levels of neonatal acute phase proteins, key components of innate immune function, measured in neonatal dried blood spots.

**Method:** We included 924 ASD cases, 1092 unaffected population-based controls, and 203 unaffected siblings to ASD cases in this case-control study nested within the register-based Stockholm Youth Cohort. Concentrations of nine different acute phase proteins were measured in eluates from neonatal dried blood spots from cases, controls, and siblings using a bead-based multiplex assay.

**Results:** C reactive protein was consistently associated with odds of ASD in case-control comparisons, with higher odds associated with the highest quintile compared to the middle quintile (OR 1.50, 95% CI 1.10 – 2.04) in adjusted analyses. In contrast, the lowest quintiles of alpha-2-macroglobulin (3.71, 1.21 – 11.33), ferritin (4.20, 1.40 – 12.65), and Serum Amyloid P (3.05, 1.16 – 8.01) were associated with odds of ASD in the matched sibling comparison. Neonatal acute phase proteins varied with perinatal environmental factors and maternal/fetal phenotypes. Significant interactions in terms of risk for ASD were observed between neonatal acute phase proteins and maternal infection in late pregnancy, maternal anemia, and maternal psychiatric history.

**Conclusions:** Indicators of the neonatal innate immune response are associated with risk for ASD, though the nature of these associations varies considerably with factors in the perinatal environment and the genetic background of the comparison group.

## Introduction

Perturbations in immune signaling pathways are hypothesized to influence neurodevelopment, as many immune effector molecules have pleiotropic effects in the developing nervous system (1, 2). Peripheral concentrations of cytokines and chemokines involved in the innate and adaptive immune responses vary in individuals already diagnosed with ASD compared to controls (3, 4), though the prevalence of multiple disorders of the immune system (e.g., asthma, infections, allergy, autoimmune disorders) is also higher in individuals with ASD (5–7). Integrative analyses of genetic and transcriptomic data suggest that ASD and several of these commonly co-occurring conditions may be related via shared mechanisms involving innate immunity (8).

In studies employing biological samples collected from etiologically-relevant periods, deviations in neonatal and fetal markers of inflammation and innate immunity exist in some individuals later diagnosed with ASD (9–14). Such differences may be attributable to genetic background or differential exposure to environmental insults already *in utero* (15–19). Acute phase proteins (APP) are key components of the innate immune response. We investigated the associations between neonatal APP and later diagnosis of ASD, and how environmental exposures (e.g., maternal infections) may affect such associations. We also compared individuals with ASD to their unaffected siblings to understand the role of shared familial factors (e.g., genetic background).

## Material and Methods

### Study population

The Stockholm Youth Cohort (SYC) is a register-based cohort of all children aged 0-17 years living in Stockholm County (20, 21). Of the individuals born 1996-2000, we selected all individuals diagnosed with ASD before 12-31-2011, a random sample of individuals in the SYC, and unaffected siblings to ASD cases (Figure S1). Diagnostic information in the SYC was updated after sample collection to include follow-up until 12-31-2016. Ethical approval was obtained the Stockholm regional review board (DNR 2010/1185-31/5). Individual consent was not required for this anonymized register-based study.

### ASD ascertainment

We implemented a case-finding procedure covering all pathways to child and adolescent psychiatric care and habilitation services in Stockholm County (20, 21). Ascertainment of ASD, ID, and ADHD in the SYC is described in Table S1. The outcome of ASD was stratified by presence of co-occurring ADHD and ID: ASD with ID, ASD with ADHD, and ASD without ID or ADHD (ASD only). Individuals diagnosed with both co-occurring ID and ADHD (n=111) were included in the ASD with ID group.

### Laboratory analysis

Neonatal dried blood spots (NDBS) were collected from the national biobank at Karolinska University Hospital, Solna. Blood spots were originally collected at approximately 3-5 days after birth (mean=4.1 days, SD=1.3). We selected a smaller sample (born 1998-2000) of the NDBS collected for analysis of immune markers. A sample (12.5 %) of those born 1996-1997 was also included. Our final sample consisted of 924 ASD cases and 1092 controls, as well as 203 unaffected siblings to ASD cases (Figure S1).

A 3.2 mm diameter disc was punched from each NDBS, immersed in 200 μL of phosphate buffered saline, and incubated at room temperature on a rotary shaker (600 rpm) for two hours. Samples for the case-cohort comparison were eluted in three batches over three subsequent days and stored at - 80°C until analysis. For the sibling comparison, a new punch was taken for each affected sibling and their unaffected sibling, placed in adjacent wells on the same plate for elution, and analyzed immediately. Total protein concentration in eluates was measured using Direct Detect infrared spectroscopy (Merck, Darmstadt, Germany).

Eluates were analyzed for procalcitonin (PCT), ferritin (FER), tissue plasminogen activator (tPA), fibrinogen (FIB), and serum amyloid A (SAA) or diluted 1:4 for analysis of α-2 macroglobulin (A2M), C reactive protein (CRP), haptoglobin (HAP), and serum amyloid P (SAP) using a premixed, multiplex panel and the Bio-Plex 200 System (Bio-Rad, Hercules, CA, USA). Coefficients of variation for manufacturer-provided controls varied by analyte from 9.0% to 20.8% (Table S2). The percentage of samples below the lower limit of quantification (LLOQ) varied by analyte from 0.00% to 0.45% (Table S2), and were assigned the value of LLOQ/√2.

### Covariates

Covariates were chosen based on previous associations with ASD. Covariate data were extracted from the Medical Birth Register, the Integrated Database for Labor Market Research, and the National Patient Register. We considered maternal age (22), country of origin (23), BMI (24), psychiatric history (25), hospitalization for infection during late pregnancy (3^rd^ trimester) (15), anemia (26), and supplement use during pregnancy (27); parental income at the time of birth and education level (highest of mother or father) (28); children’s birth order, sex, gestational age at birth (29), size for gestational age (30), mode of delivery (31), and Apgar score (32).

### Statistical analysis

We log_2_-transformed the APP values as the distributions were right skewed. Because of differences in total protein and APP distributions according to elution batch (n=3) and analytical plate (n=24; Figure S2), we created plate-specific standardized z-scores for each APP by subtracting the plate-specific mean from each observation and dividing by the plate-specific standard deviation. We created quintiles of each APP, using the distribution of z-scores among unaffected individuals to set cut-offs.

We examined the association of each covariate with the APP z-scores, separately for ASD cases and controls, estimating mean APP z-score over categories of the covariates with linear regression, followed by a Wald test as a general test of the association between the APP and the covariate.

We used conditional logistic regression models stratified by plate to calculate the odds of ASD associated with each APP. For categorical analysis, the middle quintile was the referent group. For continuous analysis, we used restricted cubic spline models with three knots. Adjusted models included terms for: maternal age, psychiatric history, country of origin, and hospitalization for infection during pregnancy; parental income; children’s birth order, sex, gestational age at birth, size for gestational age, and mode of delivery. We tested for interactions between APP levels and the covariates, comparing models including the cross-product terms to those without the cross-product terms using a likelihood ratio test. Matched sibling analysis used conditional logistic regression models, clustered by family, and adjusted for sex, birth order, and total protein concentration.

We conducted sensitivity analyses to determine whether variation in APP measurements may have influenced results. First, we additionally adjusted for factors that influence APP levels but were not associated with ASD (total protein, birth year, and age at neonatal blood sample). Second, we excluded samples with low total protein (i.e., those within the lowest quartile of total protein). Finally, we used multi-level modeling to account for both plate and batch.

## Results

### Association of APP with covariates

All APPs were correlated with each other and with total protein yield (p<0.05) (Figure S3). Other than parental education, maternal BMI, and maternal psychiatric history, each covariate was also associated with at least one APP (p<0.05; Figure 1; Table S4), though the patterns of association varied for each APP. For example, CRP was the only APP associated with sex, with a higher z-score for males (0.03) than females (−0.16; p=0.001). The patterns of association between APP and covariates differed among cases versus controls (Figure S4, Table S5). For example, levels of most APP, particularly FER, FIB, PCT, and tPA, were lower among cases born to mothers who had been hospitalized for infection during the third trimester compared to levels among cases whose mothers had not been hospitalized for infections and levels of FER were lower among cases born to mothers diagnosed with anemia (Figure S4), in contrast to the patterns observed among controls (Figure 1).

**Fig. 1.**
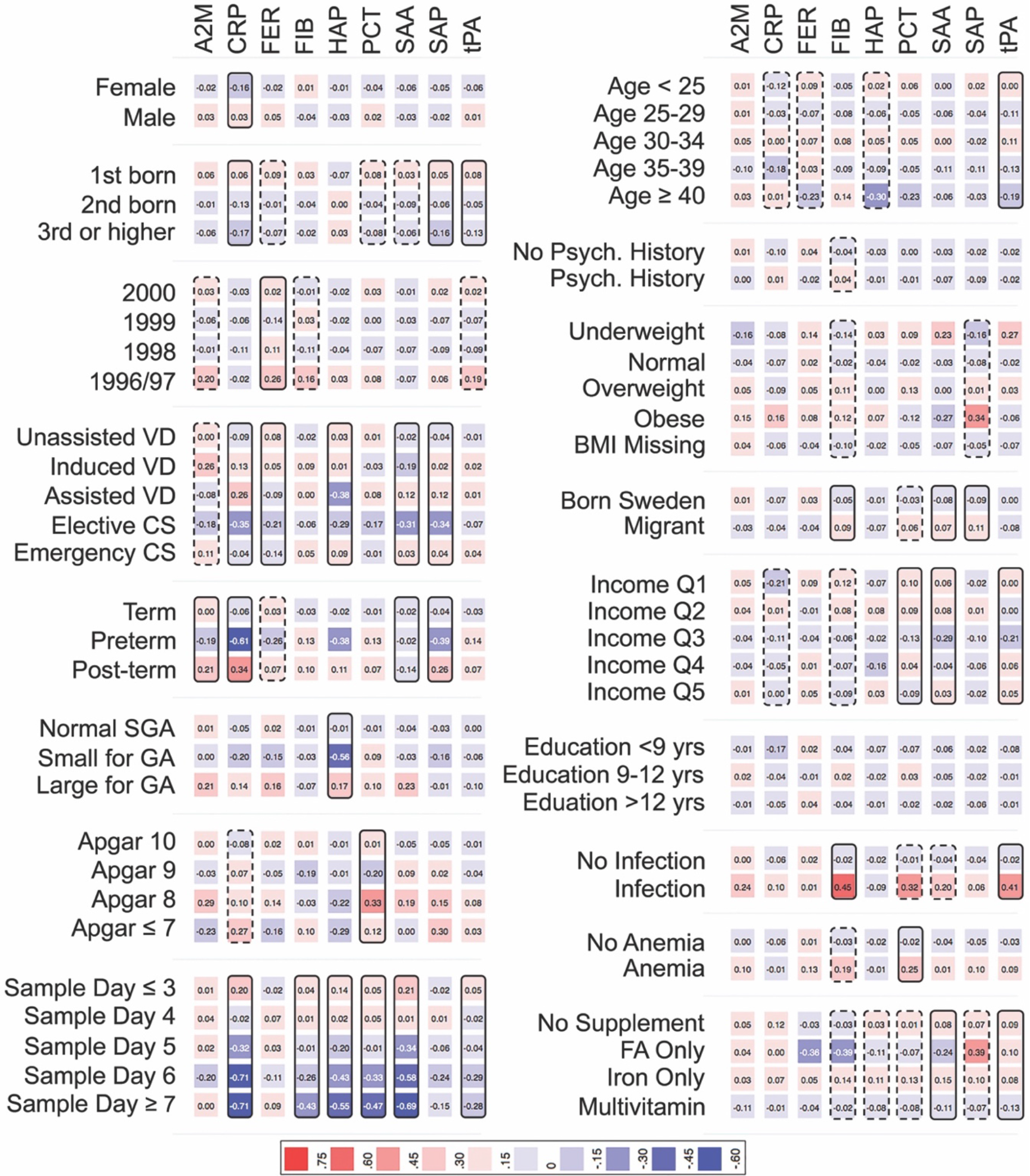
Heat map showing the mean APP z-score according to each category of the covariates. among 1092 unaffected individuals in the cohort. Solid boxes indicate that the APP is associated with the covariate at p<0.05. Dashed boxes indicate that the APP is associated with the covariate at p<0.20. Abbreviations: **A2M**: α-2 macroglobulin; **CRP**: C reactive protein; **FER**: ferritin; **FIB**: fibrinogen; **HAP**: haptoglobin; **PCT**: procalcitonin; **SAA**: serum amyloid A; **SAP**: serum amyloid P; and **tPA**: tissue plasminogen activator; **VD**: vaginal delivery; **CS**: Caesarean section delivery; **SGA**: size for gestational age; **GA**: gestational age; **Psych:** Psychiatric; **BMI**: body mass index; **Income Q**: income quintile; **FA**: folic acid.

### Association of APP with odds of ASD

Median levels of all APP except FER were higher in ASD cases compared to controls (Table 1), though the overall distributions of APP were similar between cases and controls (Figure S5). In regression analyses, we observed modest u-shaped associations between CRP, FIB, PCT, and SAA and odds of ASD, with the strongest associations between CRP and odds of ASD (Figure 2) and evidence for non-linear associations for CRP, FIB, and PCT (Table S6). The relationships between FIB, PCT, and SAA and ASD were attenuated in the fully adjusted models. Similar patterns of association were observed when we considered APP as a categorical variable, with increased odds of ASD associated with the highest quintiles of CRP (OR 1.50, 95% CI 1.10 – 2.04) and PCT (1.35, 0.99 – 1.85) compared to the middle quintile in adjusted models (Figure S6A).

**Fig. 2.**
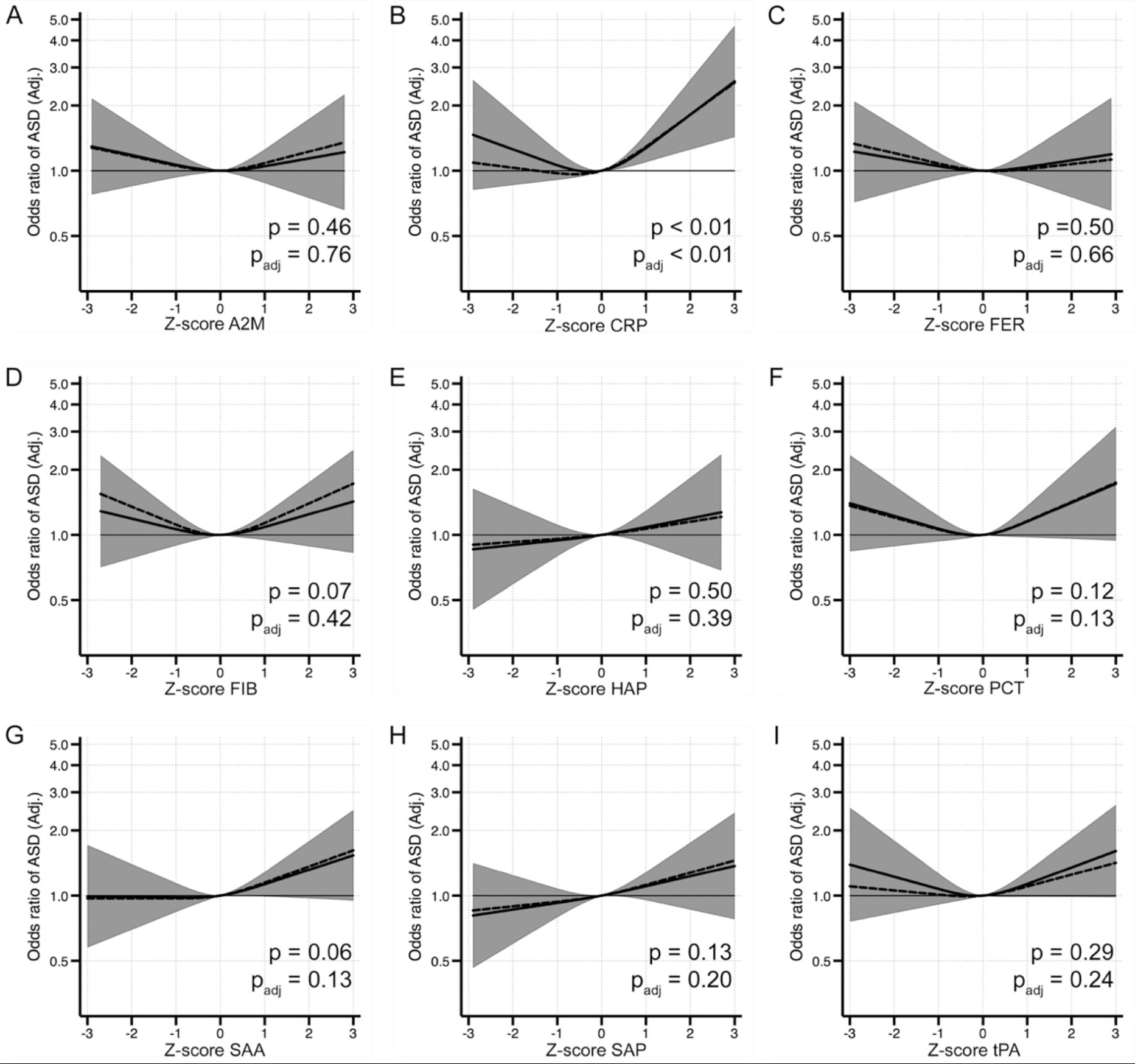
The relationship between APP and odds of ASD when comparing 924 ASD cases to 1092 unaffected individuals selected from the cohort. Each panel displays the odds of ASD according to APP z-score, flexibly fit using restricted cubic spline models with three knots and a z-score=0 as the referent. The dashed line represents the unadjusted estimate of the relationship between each APP and odds of ASD. The solid line represents the fully adjusted model, adjusted for maternal age, psychiatric history, country of origin, and hospitalization for infection during pregnancy; children’s birth order, sex, gestational age at birth, size for gestational age, and mode of delivery. The gray bands represent the 95% confidence interval for the fully adjusted model. P-values are shown for a Wald test with a null hypothesis that all APP spline terms were jointly equal to zero, as a test of whether each APP was generally associated with the outcome. P values are shown for spline terms in both the unadjusted models (p) and the adjusted models (p_adj_). Abbreviations: **ASD**: autism spectrum disorders; **A2M**: α-2 macroglobulin; **CRP**: C reactive protein; **FER**: ferritin; **FIB**: fibrinogen; **HAP**: haptoglobin; **PCT**: procalcitonin; **SAA**: serum amyloid A; **SAP**: serum amyloid P; and **tPA**: tissue plasminogen activator.

**Table 1.** Characteristics of individuals diagnosed with ASD and unaffected individuals in the study sample.

|  | Unaffected<br>(n = 1092) | ASD<br>(n = 924) | p-value<sup>a</sup> | ASD with ID<br>(n = 321) | ASD with<br>ADHD<br>(n = 322) | ASD only<br>(n = 281) | p-value<sup>b</sup> |
| --- | --- | --- | --- | --- | --- | --- | --- |
| **Sex** |  |  |  |  |  |  |  |
| Female | 538 (49.3%) | 214 (23.2%) | <0.001 | 84 (26.2%) | 70 (21.7%) | 60 (21.4%) | <0.001 |
| Male | 554 (50.7%) | 710 (76.8%) |  | 237 (73.8%) | 252 (78.3%) | 221 (78.6%) |  |
| **Birth Order** |  |  |  |  |  |  |  |
| 1st born | 443 (40.6%) | 419 (45.3%) | 0.030 | 124 (38.6%) | 158 (49.1%) | 137 (48.8%) | 0.006 |
| 2nd born | 385 (35.3%) | 289 (31.3%) |  | 114 (35.5%) | 87 (27.0%) | 88 (31.3%) |  |
| 3rd or higher | 205 (18.8%) | 148 (16.0%) |  | 60 (18.7%) | 49 (15.2%) | 39 (13.9%) |  |
| Missing | 59 (5.4%) | 68 (7.4%) |  | 23 (7.2%) | 28 (8.7%) | 17 (6.0%) |  |
| **Mode of Delivery** |  |  |  |  |  |  |  |
| Unassisted VD | 712 (65.2%) | 532 (57.6%) | 0.014 | 175 (54.5%) | 180 (55.9%) | 177 (63.0%) | 0.042 |
| Induced VD | 63 (5.8%) | 71 (7.7%) |  | 23 (7.2%) | 30 (9.3%) | 18 (6.4%) |  |
| Assisted VD | 102 (9.3%) | 90 (9.7%) |  | 37 (11.5%) | 29 (9.0%) | 24 (8.5%) |  |
| Elective CS | 65 (6.0%) | 71 (7.7%) |  | 26 (8.1%) | 21 (6.5%) | 24 (8.5%) |  |
| Emergency CS | 91 (8.3%) | 92 (10.0%) |  | 37 (11.5%) | 34 (10.6%) | 21 (7.5%) |  |
| Missing | 59 (5.4%) | 68 (7.4%) |  | 23 (7.2%) | 28 (8.7%) | 17 (6.0%) |  |
| **Gestational Age** |  |  |  |  |  |  |  |
| ≤36 weeks | 53 (4.9%) | 63 (6.8%) | 0.12 | 23 (7.2%) | 21 (6.5%) | 19 (6.8%) | 0.59 |
| 37-41 weeks | 886 (81.1%) | 723 (78.2%) |  | 249 (77.6%) | 250 (77.6%) | 224 (79.7%) |  |
| ≥42 weeks | 92 (8.4%) | 69 (7.5%) |  | 26 (8.1%) | 23 (7.1%) | 20 (7.1%) |  |
| Missing | 61 (5.6%) | 69 (7.5%) |  | 23 (7.2%) | 28 (8.7%) | 18 (6.4%) |  |
| **Size for Gestational Age** |  |  |  |  |  |  |  |
| Normal size for GA | 960 (87.9%) | 772 (83.5%) | 0.17 | 274 (85.4%) | 258 (80.1%) | 240 (85.4%) | 0.032 |
| Small for GA | 29 (2.7%) | 29 (3.1%) |  | 14 (4.4%) | 10 (3.1%) | 5 (1.8%) |  |
| Large for GA | 36 (3.3%) | 43 (4.7%) |  | 8 (2.5%) | 20 (6.2%) | 15 (5.3%) |  |
| Missing | 67 (6.1%) | 80 (8.7%) |  | 25 (7.8%) | 34 (10.6%) | 21 (7.5%) |  |
| **Apgar Score at 5 Minutes** |  |  |  |  |  |  |  |
| 10 | 861 (78.8%) | 685 (74.1%) | 0.064 | 219 (68.2%) | 243 (75.5%) | 223 (79.4%) | <0.001 |
| 9 | 110 (10.1%) | 101 (10.9%) |  | 35 (10.9%) | 36 (11.2%) | 30 (10.7%) |  |
| 8 | 33 (3.0%) | 36 (3.9%) |  | 20 (6.2%) | 10 (3.1%) | 6 (2.1%) |  |
| ≤7 | 15 (1.4%) | 25 (2.7%) |  | 19 (5.9%) | 3 (0.9%) | 3 (1.1%) |  |
| Missing | 73 (6.7%) | 77 (8.3%) |  | 28 (8.7%) | 30 (9.3%) | 19 (6.8%) |  |
| **Age at Neonatal Blood Sample (Days)** |  |  |  |  |  |  |  |
| ≤3 | 423 (38.7%) | 346 (37.4%) | 0.19 | 109 (34.0%) | 127 (39.4%) | 110 (39.1%) | 0.13 |
| 4 | 357 (32.7%) | 312 (33.8%) |  | 106 (33.0%) | 114 (35.4%) | 92 (32.7%) |  |
| 5 | 195 (17.9%) | 167 (18.1%) |  | 68 (21.2%) | 51 (15.8%) | 48 (17.1%) |  |
| 6 | 84 (7.7%) | 55 (6.0%) |  | 17 (5.3%) | 19 (5.9%) | 19 (6.8%) |  |
| ≥7 | 31 (2.8%) | 41 (4.4%) |  | 21 (6.5%) | 10 (3.1%) | 10 (3.6%) |  |
| Missing | 2 (0.2%) | 3 (0.3%) |  | 0 (0.0%) | 1 (0.3%) | 2 (0.7%) |  |
| **Maternal Age (Years)** |  |  |  |  |  |  |  |
| >25 | 145 (13.3%) | 116 (12.6%) | 0.011 | 51 (15.9%) | 40 (12.4%) | 25 (8.9%) | <0.001 |
| 25-29 | 293 (26.8%) | 307 (33.2%) |  | 108 (33.6%) | 118 (36.6%) | 81 (28.8%) |  |
| 30-34 | 393 (36.0%) | 291 (31.5%) |  | 86 (26.8%) | 102 (31.7%) | 103 (36.7%) |  |
| 35-39 | 229 (21.0%) | 173 (18.7%) |  | 56 (17.4%) | 56 (17.4%) | 61 (21.7%) |  |
| ≥40 | 32 (2.9%) | 37 (4.0%) |  | 20 (6.2%) | 6 (1.9%) | 11 (3.9%) |  |
| **Maternal Psychiatric History** |  |  |  |  |  |  |  |
| No | 723 (66.2%) | 483 (52.3%) | <0.001 | 190 (59.2%) | 145 (45.0%) | 148 (52.7%) | <0.001 |
| Yes | 369 (33.8%) | 441 (47.7%) |  | 131 (40.8%) | 177 (55.0%) | 133 (47.3%) |  |
| **Maternal BMI** |  |  |  |  |  |  |  |
| Normal | 538 (49.3%) | 364 (39.4%) | <0.001 | 130 (40.5%) | 110 (34.2%) | 124 (44.1%) | <0.001 |
| Underweight | 22 (2.0%) | 22 (2.4%) |  | 8 (2.5%) | 10 (3.1%) | 4 (1.4%) |  |
| Overweight | 184 (16.8%) | 165 (17.9%) |  | 61 (19.0%) | 62 (19.3%) | 42 (14.9%) |  |
| Obese | 49 (4.5%) | 77 (8.3%) |  | 25 (7.8%) | 35 (10.9%) | 17 (6.0%) |  |
| Missing | 299 (27.4%) | 296 (32.0%) |  | 97 (30.2%) | 105 (32.6%) | 94 (33.5%) |  |
| **Mother Born Outside Sweden** |  |  |  |  |  |  |  |
| No | 807 (73.9%) | 647 (70.0%) | 0.053 | 168 (52.3%) | 262 (81.4%) | 217 (77.2%) | <0.001 |
| Yes | 285 (26.1%) | 277 (30.0%) |  | 153 (47.7%) | 60 (18.6%) | 64 (22.8%) |  |
| **Family Income Quintile** |  |  |  |  |  |  |  |
| 1<sup>st</sup> (Lowest) | 154 (14.1%) | 122 (13.2%) | 0.003 | 64 (19.9%) | 29 (9.0%) | 29 (10.3%) | <0.001 |
| 2<sup>nd</sup> | 234 (21.4%) | 240 (26.0%) |  | 92 (28.7%) | 82 (25.5%) | 66 (23.5%) |  |
| 3<sup>rd</sup> | 240 (22.0%) | 214 (23.2%) |  | 68 (21.2%) | 84 (26.1%) | 62 (22.1%) |  |
| 4<sup>th</sup> | 229 (21.0%) | 205 (22.2%) |  | 64 (19.9%) | 82 (25.5%) | 59 (21.0%) |  |
| 5<sup>th</sup> | 235 (21.5%) | 141 (15.3%) |  | 32 (10.0%) | 45 (14.0%) | 64 (22.8%) |  |
| Missing | 0 (0.0%) | 2 (0.2%) |  | 1 (0.3%) | 0 (0.0%) | 1 (0.4%) |  |
| **Parental Education** |  |  |  |  |  |  |  |
| <9 years | 165 (15.1%) | 167 (18.1%) | 0.078 | 74 (23.1%) | 59 (18.3%) | 34 (12.1%) | <0.001 |
| 9-12 years | 474 (43.4%) | 411 (44.5%) |  | 140 (43.6%) | 155 (48.1%) | 116 (41.3%) |  |
| >12 years | 453 (41.5%) | 344 (37.2%) |  | 106 (33.0%) | 108 (33.5%) | 130 (46.3%) |  |
| Missing | 0 (0.0%) | 2 (0.2%) |  | 1 (0.3%) | 0 (0.0%) | 1 (0.4%) |  |
| **Maternal Hospitalization for Infection (3rd trimester)** |  |  |  |  |  |  |  |
| No | 1000 (91.6%) | 811 (87.8%) | 0.024 | 279 (86.9%) | 283 (87.9%) | 249 (88.6%) | 0.050 |
| Yes | 31 (2.8%) | 43 (4.7%) |  | 19 (5.9%) | 11 (3.4%) | 13 (4.6%) |  |
| Missing | 61 (5.6%) | 70 (7.6%) |  | 23 (7.2%) | 28 (8.7%) | 19 (6.8%) |  |
| **Maternal Anemia During Pregnancy** |  |  |  |  |  |  |  |
| No | 1030 (94.3%) | 866 (93.7%) | 0.57 | 298 (92.8%) | 302 (93.8%) | 266 (94.7%) | 0.75 |
| Yes | 62 (5.7%) | 58 (6.3%) |  | 23 (7.2%) | 20 (6.2%) | 15 (5.3%) |  |
| **Maternal Dietary Supplementation** |  |  |  |  |  |  |  |
| None | 433 (39.7%) | 383 (41.5%) | 0.23 | 119 (37.1%) | 137 (42.5%) | 127 (45.2%) | 0.39 |
| Folic Acid only | 10 (0.9%) | 14 (1.5%) |  | 4 (1.2%) | 6 (1.9%) | 4 (1.4%) |  |
| Iron only | 336 (30.8%) | 294 (31.8%) |  | 109 (34.0%) | 102 (31.7%) | 83 (29.5%) |  |
| Multivitamin | 313 (28.7%) | 233 (25.2%) |  | 89 (27.7%) | 77 (23.9%) | 67 (23.8%) |  |
| **Proteins measured in NDBS (median [IQR])** |  |  |  |  |  |  |  |
| A2M (ng/ml) | 533.0 (252.3, 911.3) | 563.3 (266.0, 1052.0) | 0.017 | 574.4 (249.7, 1051.9) | 596.0 (307.1, 1123.1) | 513.6 (251.4, 1004.0) | 0.037 |
| CRP (ng/ml) | 1.3 (0.6, 3.1) | 1.6 (0.7, 4.1) | <0.001 | 1.5 (0.7, 4.1) | 1.7 (0.6, 4.1) | 1.7 (0.7, 4.2) | 0.006 |
| FER (ng/ml) | 1.8 (0.7, 3.3) | 1.7 (0.6, 3.9) | 0.27 | 1.7 (5.9, 3.6) | 2.2 (0.8, 4.2) | 1.5 (0.5, 4.0) | 0.067 |
| FIB (ng/ml) | 27.1 (16.5, 53.1) | 30.6 (16.5, 60.6) | 0.074 | 30.7 (16.9, 63.1) | 32.1 (17.5, 61.1) | 27.1 (15.4, 55.9) | 0.098 |
| HAP (ng/ml) | 27.5 (9.3, 76.8) | 29.9 (11.0, 85.4) | 0.036 | 27.6 (10.6, 78.2) | 33.7 (12.1, 87.9) | 29.1 (9.8, 88.5) | 0.10 |
| PCT (pg/ml) | 10.9 (7.4, 15.6) | 11.7 (7.5, 17.1) | 0.013 | 11.0 (7.7, 16.9) | 12.0 (7.7, 17.9) | 11.1 (7.0, 17.6) | 0.038 |
| SAA (ng/ml) | 1.3 (0.8, 2.2) | 1.4 (0.9, 2.5) | 0.019 | 1.4 (0.8, 2.7) | 1.5 (0.9, 2.7) | 1.3 (0.8, 2.2) | 0.030 |
| SAP (ng/ml) | 10.5 (6.8, 15.5) | 11.5 (7.5, 17.7) | <0.001 | 11.1 (8.1, 18.4) | 12.0 (7.5, 17.5) | 11.2 (7.0, 16.9) | 0.007 |
| tPA (pg/ml) | 9.6 (6.2, 14.7) | 10.6 (6.6, 15.9) | 0.010 | 9.9 (6.6, 15.0) | 11.5 (7.2, 17.4) | 10.1 (6.3, 16.1) | 0.004 |
| Total protein (mg/ml) | 1.3 (1.0, 1.7) | 1.3 (1.0, 1.7) | 0.76 | 1.2 (0.9, 1.6) | 1.3 (1.0, 1.7) | 1.3 (1.0, 1.7) | 0.19 |

- Pearson's chi-squared test was used for categorical variables, comparing the frequency distributions among unaffected individuals to the distributions among all ASD-affected individuals. Kruskal-Wallis tests were used for continuous variables, as the distributions of the APP concentrations were strongly skewed. - Pearson's chi-squared test was used for categorical variables, comparing the frequency distributions among unaffected individuals to the distribution among the stratified ASD outcome groups. Kruskal-Wallis tests were used for continuous variables, as the distributions of the APP concentrations were strongly skewed.
Abbreviations: **ASD**: autism spectrum disorders; **ADHD**: attention-deficit/hyperactivity disorder; **ID**: intellectual disability; **VD**: vaginal delivery; **CS**: Caesarean section delivery; **GA**: gestational age; **BMI**: body mass index; **IQR**: interquartile range; **A2M**: α-2 macroglobulin; **CRP**: C reactive protein; **FER**: ferritin; **FIB**: fibrinogen; **HAP**: haptoglobin; **PCT**: procalcitonin; **SAA**: serum amyloid A; **SAP**: serum amyloid P; and **tPA**: tissue plasminogen activator.

Overall the patterns of association with APP were similar among the stratified outcomes. The odds associated with the highest quintile of CRP were greatest for ASD only (1.80, 1.15–2.82) compared to ASD with co-occurring ID (1.29, 0.83 – 2.01) or with co-occurring ADHD (1.34, 0.85–2.11), with similar results in continuous analyses (Figures S7–10).

### APP and Odds of ASD Among Matched Sibling Pairs

In contrast to the case-control comparison, we found that median levels of all APP except CRP, HAP, and PCT were lower in ASD cases compared to their unaffected siblings (Table S3), with distinct distributions of particularly A2M, SAP, and FER among ASD cases compared to their unaffected siblings (Figure S5). In matched regression analyses, we observed decreasing odds with increasing levels of A2M, FER, SAA, SAP, and tPA (Figure 3, Figure S6B). Associations with SAA, SAP, and tPA were attenuated in adjusted models, though the pattern associated with A2M and FER persisted (Figure 3). Compared to the middle quintile, the lowest quintile of A2M (3.71, 1.21 – 11.33), FER (4.20, 1.40 – 12.65), and SAP 3.05 (1.16 – 8.01) were associated with increased odds of ASD in the matched sibling comparison (Figure S6B).

**Fig. 3.**
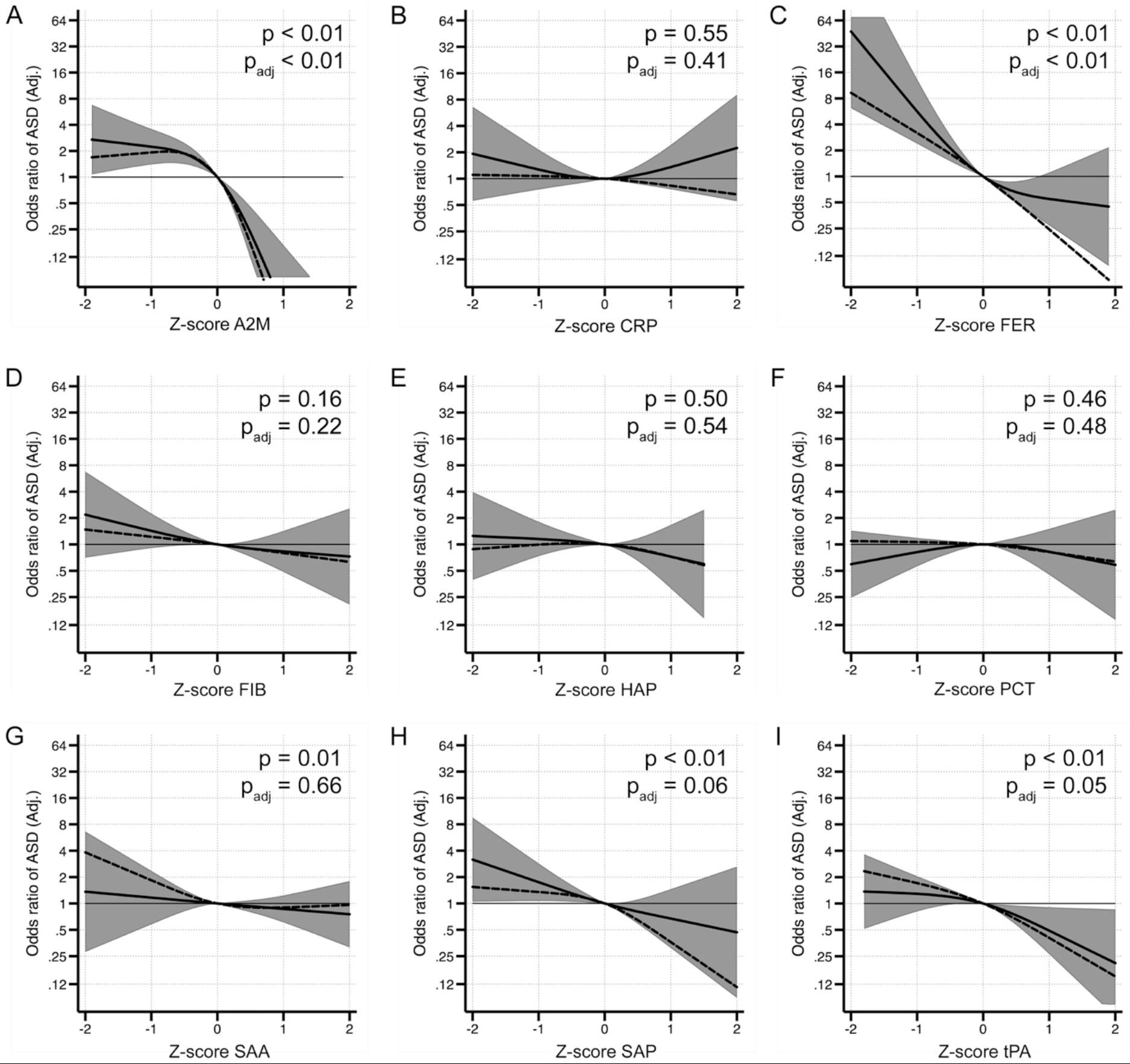
The relationship between APP and odds of ASD when comparing 203 ASD cases to their matched, unaffected sibling. Each panel displays the odds of ASD according to APP z-score, flexibly fit using restricted cubic spline models with three knots and a z-score=0 as the referent. The dashed line represents the unadjusted estimate of the relationship between each APP and odds of ASD. The solid line represents the adjusted model, adjusted for children’s birth order, and the total protein concentration of the sample. The gray bands represent the 95% confidence interval for the fully adjusted model. P-values are shown for a Wald test with a null hypothesis that all APP spline terms were jointly equal to zero, as a test of whether each APP was generally associated with the outcome. P values are shown for spline terms in both the unadjusted models (p) and the adjusted models (p_adj_). Abbreviations: **ASD**: autism spectrum disorders; **A2M**: α-2 macroglobulin; **CRP**: C reactive protein; **FER**: ferritin; **FIB**: fibrinogen; **HAP**: haptoglobin; **PCT**: procalcitonin; **SAA**: serum amyloid A; **SAP**: serum amyloid P; and **tPA**: tissue plasminogen activator.

### Interaction Between APP and Other Risk Factors for ASD

Since many of the covariates we studied (e.g., maternal infection, mode of delivery) represent environmental influences that may influence the innate immune system and are also risk factors for ASD, we hypothesized that variation in the innate immune response to these environmental influences may be associated with risk for ASD. We observed interactions between APP and a number of other risk factors for ASD in our study (Figure 4, Figure S11, Table S7). We observed interactions between maternal hospitalization for infections during the third trimester and neonatal levels of APP (Figure 4). Lower odds of ASD were associated with increasing levels of A2M, FER, FIB, PCT, and tPA among children whose mothers were hospitalized for infection late in pregnancy (Figure 4A-E), in contrast to those whose mothers were not hospitalized for infection. Similarly, we observed lower odds of ASD associated with increasing levels of FER and SAP among children whose mothers were diagnosed with anemia during pregnancy, in contrast to children whose mothers were not diagnosed with anemia (Figure 4F-G). The relationship between elevated CRP and ASD was stronger among children whose mother had no history of psychiatric illness compared to those whose mother did have a psychiatric history (Figure 4H).

**Fig. 4.**
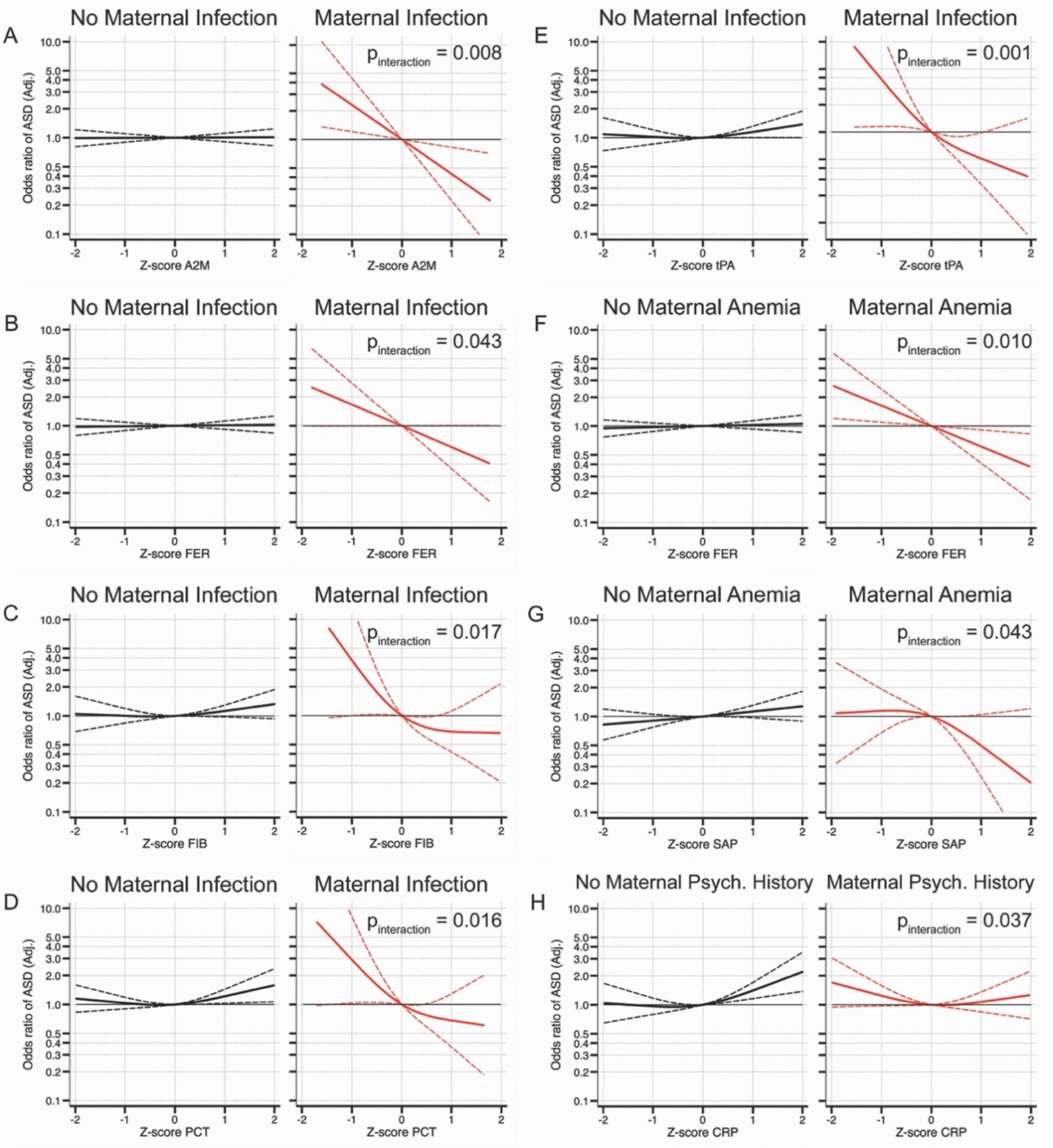
Interaction between APP and other risk factors. Odds ratios for ASD over the range of APP are shown separately for those who were not exposed to maternal hospitalization for infection in late pregnancy (3^rd^ trimester) compared to those who were (A-E), those who were not exposed to maternal anemia compared to those who were (F-G), and those whose mother had no history of psychiatric diagnoses compared to those whose mother did have a psychiatric history (H). Solid lines represent the OR estimate for each group and dashed lines represent the 95% confidence interval for the fully adjusted model. P-values for interaction (a likelihood ratio test comparing a model with interaction terms to a model without interaction terms) are shown.

### Sensitivity analyses

Sensitivity analyses were designed to evaluate whether variation in APP measurements may have influenced results. Estimates from models in sensitivity analyses were similar to the results of the main analysis (Figure S12).

## Discussion

Our results indicate that of the innate immune markers that we studied, only CRP was associated with risk for ASD, with levels of CRP above the population mean associated with increased odds of ASD even after accounting for a range for potential confounders. We observed significant interactions between environmental exposures that could plausibly influence the neonatal innate immune system and levels of APP. With the exception of PCT, higher levels of these markers also tended to be associated with lower risk of ASD when comparing ASD cases to their unaffected siblings.

### Strengths and Weaknesses

The setting of the study within a universal healthcare system with developmental screening provided to families free of charge increases the likelihood that cases of ASD are identified, using validated ascertainment methods (21). We included two comparison groups in the study. Unaffected siblings share aspects of the early life environment with ASD-affected siblings as well as on average 50% of their common genetic variants. The large population-based comparison group of unrelated individuals randomly sampled from the larger cohort included in the study is also a strength, allowing us to study the relationship between a large number of potential confounders and neonatal levels of APP. While the use of multiple markers of the innate immune response and information on many putative risk factors for ASD is a strength of this study, there is still a possibility for residual confounding and the analysis of such markers results in multiple statistical comparisons.

The use of APP as markers of the neonatal innate immune response is also a strength of this study. Previous studies have measured cytokines and chemokines, which is technically challenging (9–12, 33). Cytokines and chemokines have short half-lives relative to acute phase proteins (34, 35). Therefore, it is difficult to interpret the “cytokine profile” with little information about the environmental factors that may influence such levels, with some studies indicating opposing effects of the same cytokines (10, 11, 33).

### Interpretation

APP are a diverse group of proteins that collectively recognize and opsonize invading pathogens and cellular debris, regulate blood viscosity and clotting, and sequester nutrients (e.g., iron) from pathogens (36). Moreover, many of the APP play key roles in the regulation of the innate and adaptive immune responses and are required for resolution of an acute inflammatory response in experimental models (37–40). Expression of APP occurs primarily in the liver, is influenced by genetic background, and increases by orders of magnitude in response to IL-6 and other inflammatory cytokines (41–44). Fetal liver produces APP, with expression detectable from early in the second trimester (45). APP are not believed to cross the placenta (46). We consider APP in NDBS as specific indicators of the neonatal innate immune status.

CRP is a pattern recognition molecule, opsonin, and activator of the complement pathway (39). Of the APP studied, CRP had the strongest association with risk of ASD in the case-control comparison, displaying a U-shaped association with odds of ASD. While the shape of the association between CRP and odds of ASD in the sibling comparison was similar, the association was less apparent with considerably wider confidence intervals in the sibling analysis, possibly reflecting shared common genetic variation in innate immune signaling and risk of ASD (8). We also observed a weaker relationship between CRP and risk of ASD among those whose mothers had a previous history of psychiatric illness in the case-control comparison, with a significant interaction detected between maternal psychiatric history and levels of CRP. Environmental stimuli that increase CRP production may be more important to the etiology of ASD among individuals with lower genetic liability for the disorder. Moreover, both positive and negative genetic correlations between psychiatric disorders (e.g., bipolar disorder, schizophrenia) and CRP levels have been reported (47, 48). Thus, levels of CRP in children born to mothers with a high genetic risk for psychiatric illness may exhibit larger variation than children to unaffected mothers.

Levels of multiple APP were elevated among neonates whose mothers were hospitalized for infections in late pregnancy, although these markers were not elevated to the same extent among children later diagnosed with ASD. In interaction analyses, higher levels of A2M, FER, FIB, PCT, and tPA were associated with lower risk of ASD among those exposed to maternal infections in late pregnancy. The actions of the APP indicated here are diverse: A2M, FIB, and tPA regulate blood coagulation, while FER binds to iron to limit the growth of pathogens. The physiological function of PCT is poorly defined, but its rapid rise in response to bacterial infections makes PCT useful when diagnosing sepsis, particularly in neonates (49). We previously reported that low levels of APP in combination with maternal exposure to cytomegalovirus and *Toxoplasma gondii* were associated with greater risk for non-affective psychosis compared to those with higher APP levels (50). Taken together with the results of the current study, this suggests that the degree of protection a neonate’s innate immune system offers against an environmental insult may be important in modulating the influence of such insults on neurodevelopmental processes. Of the APP that interacted with infection in terms of risk for ASD, higher levels of all but PCT were also associated with lower odds of ASD in the sibling comparison study. In other words, we observe that, within matched pairs, the sibling who produced higher levels of these APP in response to a presumably similar early life environment was less likely to develop ASD.

While FER concentrations rise to withhold iron from pathogens during the acute phase response (36), in the absence of this response, FER reflects the body’s store of iron in both adults and neonates (51). We recently reported that maternal diagnosis with anemia was associated with ASD, ID, and ADHD in offspring (26). The results of that study indicated that the increased risk associated with maternal anemia may be limited to cases where anemia was severe and long lasting. Here we report evidence that among individuals whose mothers were affected by anemia during pregnancy, higher levels of ferritin were associated with lower odds of ASD. These results suggest that among mothers affected by anemia, an adequate iron store in the fetus may be protective with regard to risk of ASD. We also observed a similar protective association with increasing levels of ferritin when comparing ASD cases to their unaffected siblings, in line with sibling comparisons of exposure to maternal anemia (26).

Numerous studies have posited adverse effects of intrauterine inflammatory signals to explain the associations consistently observed between elevated maternal BMI and risk for ASD (52–55). Here we report that maternal BMI was not associated with APPs in cases or controls measured only a few days after birth, raising the question of whether this particular mechanism to link maternal BMI to risk of ASD is relevant, at least in later pregnancy.

## Conclusion

Together, our findings suggest that it is not the levels of inflammatory or regulatory markers *per se* that may influence ASD risk. Rather, the strength of those signals in the genetic and environmental context of the developing nervous system must be considered. Correspondingly, understanding on a molecular level the individual’s ability to resolve environmental insults may help to identify those that are particularly susceptible to such insults.

## Ackowledgements

This work was supported by grants from the Swedish Research Council (grant numbers 2016-01477, 2012-2264 and 523-2010-1052 [to CD]), Autism Speaks (Basic and Clinical Grant #7618 [to BKL]), and the Stanley Medical Research Institute (to HK). The funders had no role in the design and conduct of the study; collection, management, analysis, and interpretation of the data; preparation, review, or approval of the manuscript; or decision to submit the manuscript for publication.

## Supplementary Materials

### Supplemental Figures

**Figure S1.**
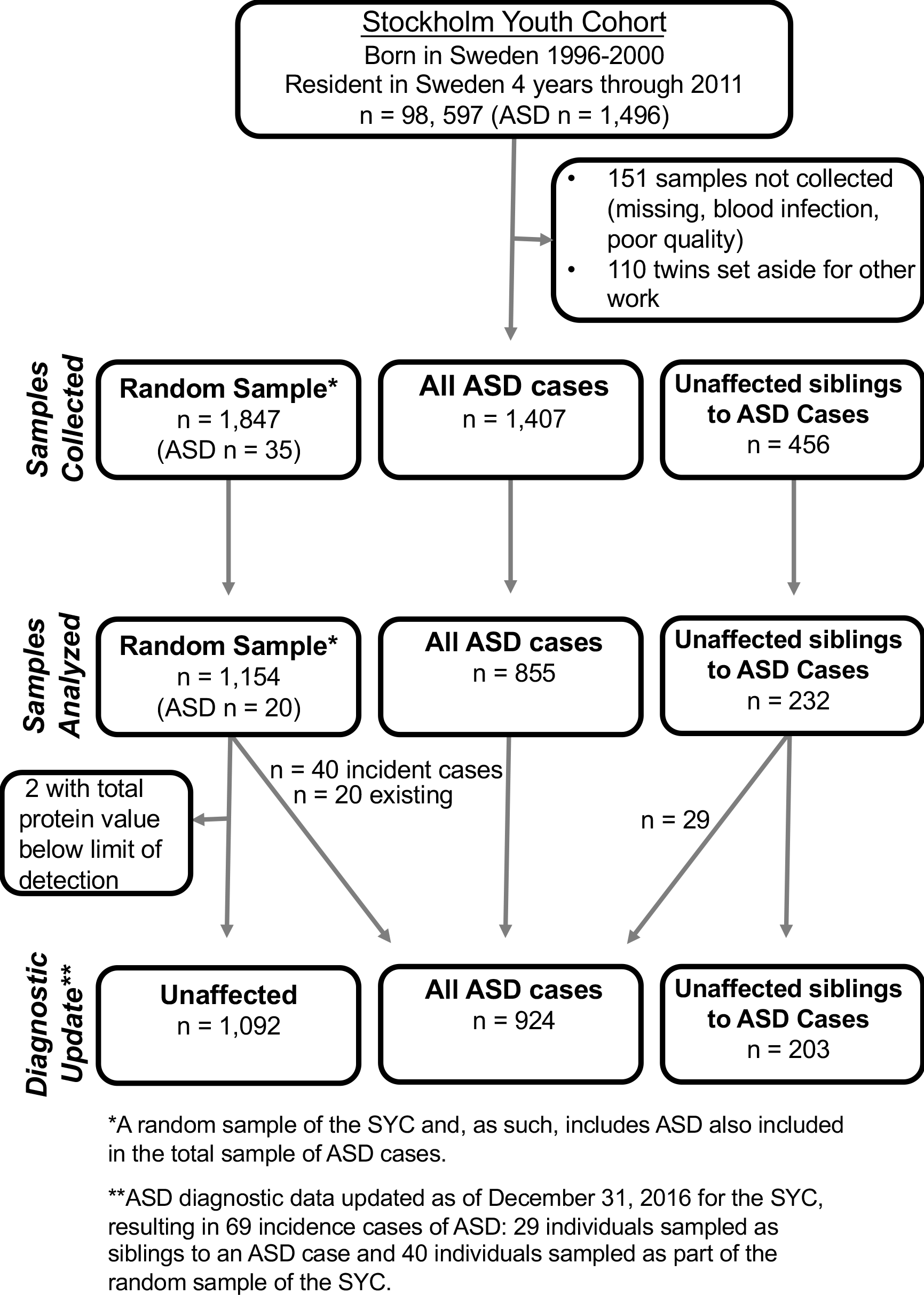
Selection of individuals and NDBS samples.

**Figure S2.**
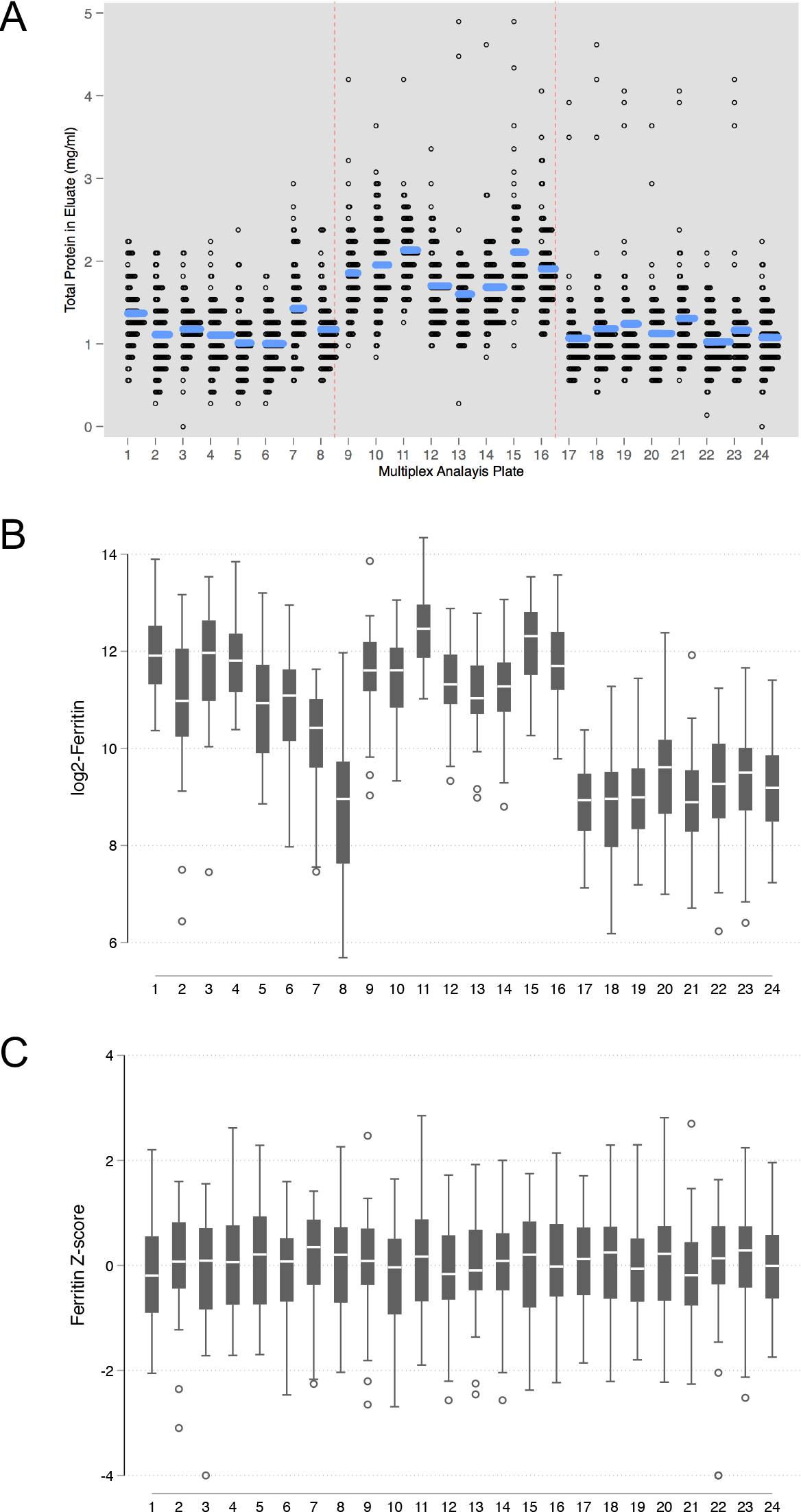
Levels of total protein (**A**) and one APP (**B** Ferritin) according to analytic plate and elution batch. Protein elution batches are demarcated by red dotted lines in **A**. Plate-specific z-scores were calculated to allow comparisons across batches and plates (**C**).

**Figure S3.**
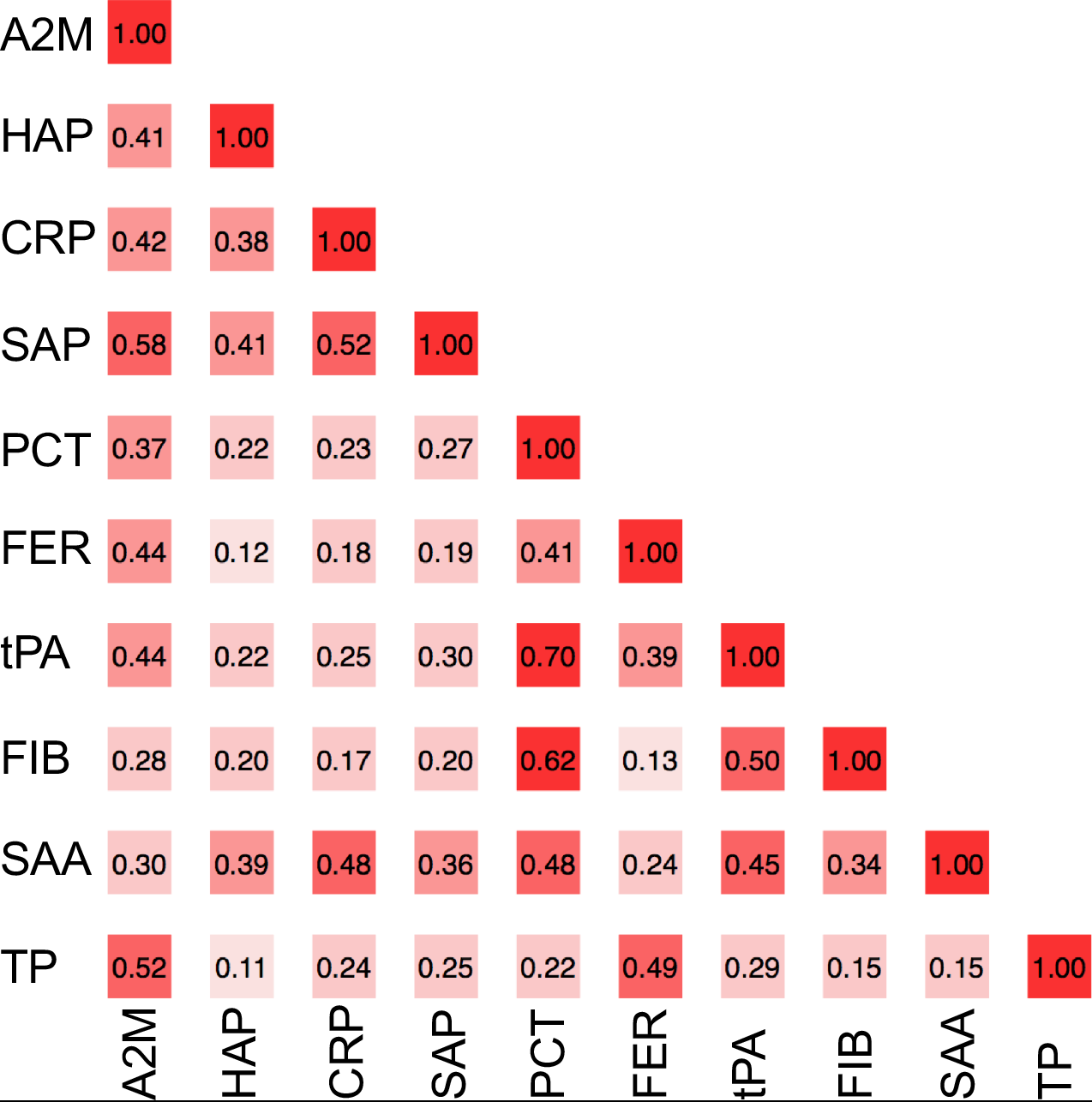
The Spearman rho correlations of nine acute phase proteins with each other and with total protein measurements, as measured in neonatal dried blood spots of 1092 unaffected controls selected from the cohort. Abbreviations: **A2M**: α-2 macroglobulin; **CRP**: C reactive protein; **FER**: ferritin; **FIB**: fibrinogen; **HAP**: haptoglobin; **PCT**: procalcitonin; **SAA**: serum amyloid A; **SAP**: serum amyloid P; **tPA**: tissue plasminogen activator; and **TP**: total protein.

**Figure S4.**
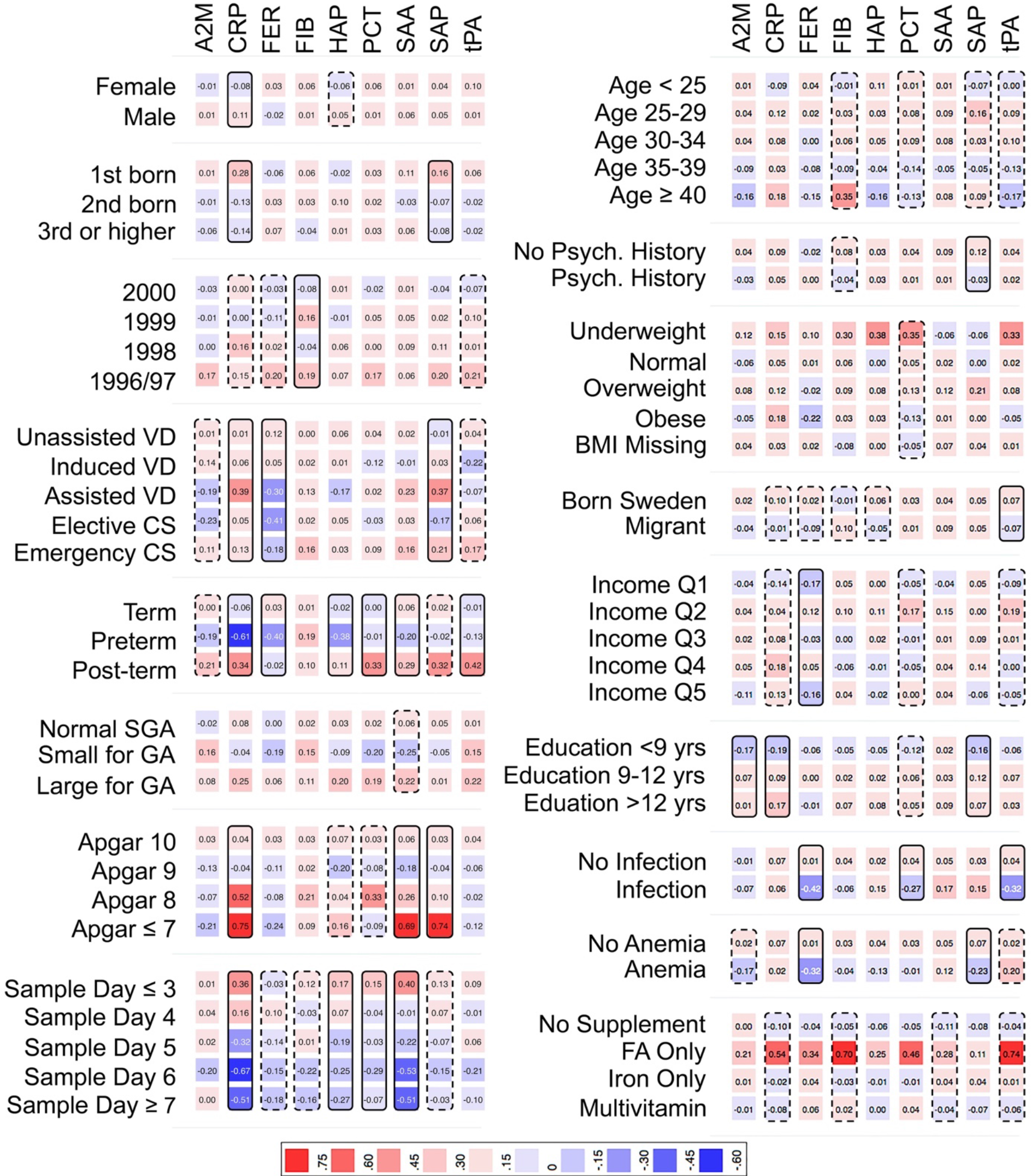
Heat map showing the mean APP z-score according to each category of the covariates, among 924 ASD-affected individuals in the cohort. Solid boxes indicate that the APP is associated with the covariate at p<0.05. Dashed boxes indicate that the APP is associated with the covariate at p<0.20. Abbreviations: **A2M**: α-2 macroglobulin; **CRP**: C reactive protein; **FER**: ferritin; **FIB**: fibrinogen; **HAP**: haptoglobin; **PCT**: procalcitonin; **SAA**: serum amyloid A; **SAP**: serum amyloid P; and **tPA**: tissue plasminogen activator; **VD**: vaginal delivery; **CS**: Caesarean section delivery; **SGA**: size for gestational age; **GA**: gestational age; **Psych:** Psychiatric; **BMI**: body mass index; **Income Q**: income quintile; **FA**: folic acid.

**Figure S5.**
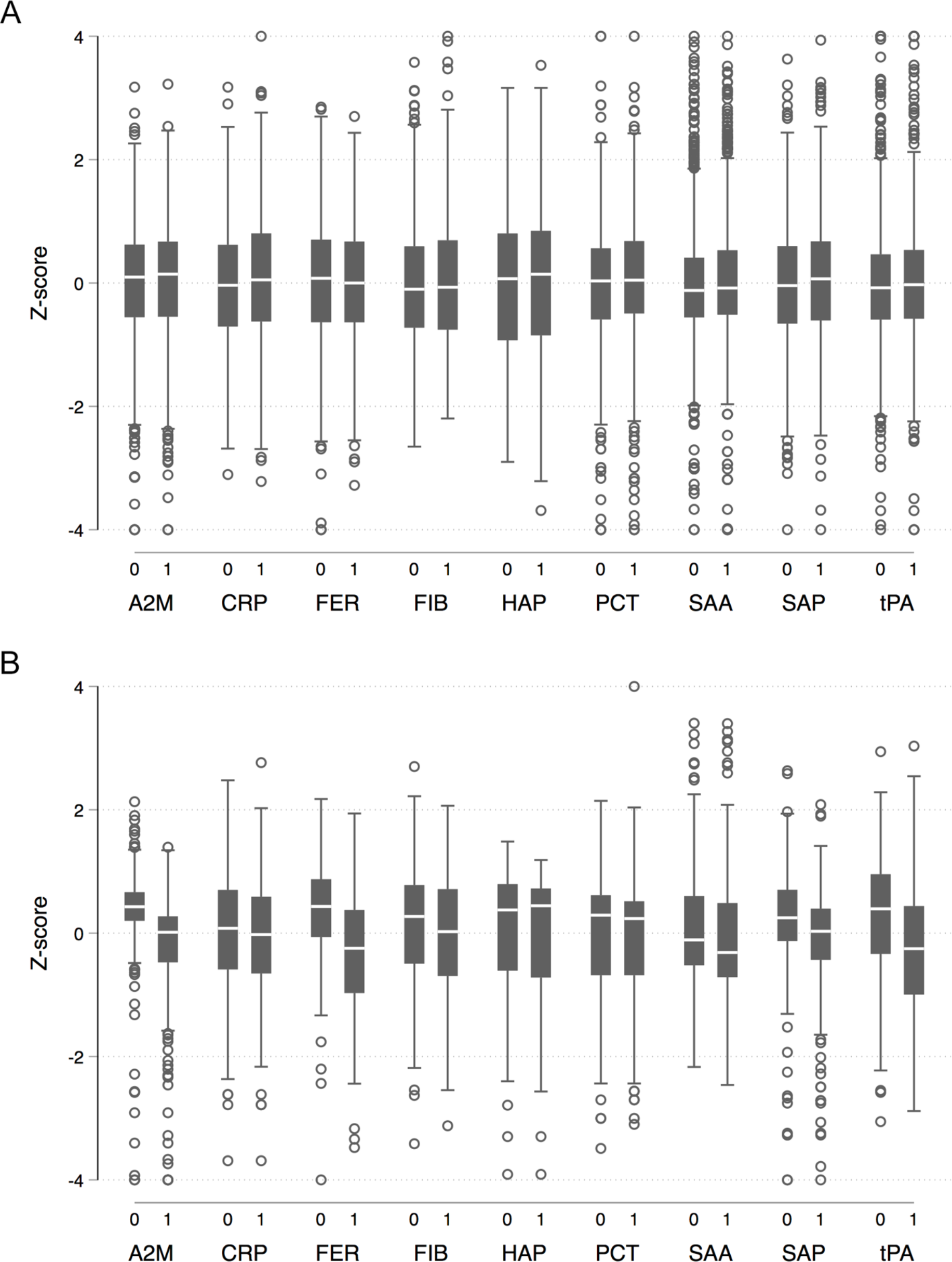
The distribution of each APP measured in neonatal dried blood spot samples, comparing ASD-unaffected individuals (0) to ASD-affected individuals (1). In (**A**), 924 ASD cases are compared to 1092 unaffected individuals selected from the cohort. In (**B**), 203 ASD cases are compared to their unaffected siblings.

**Figure S6.**
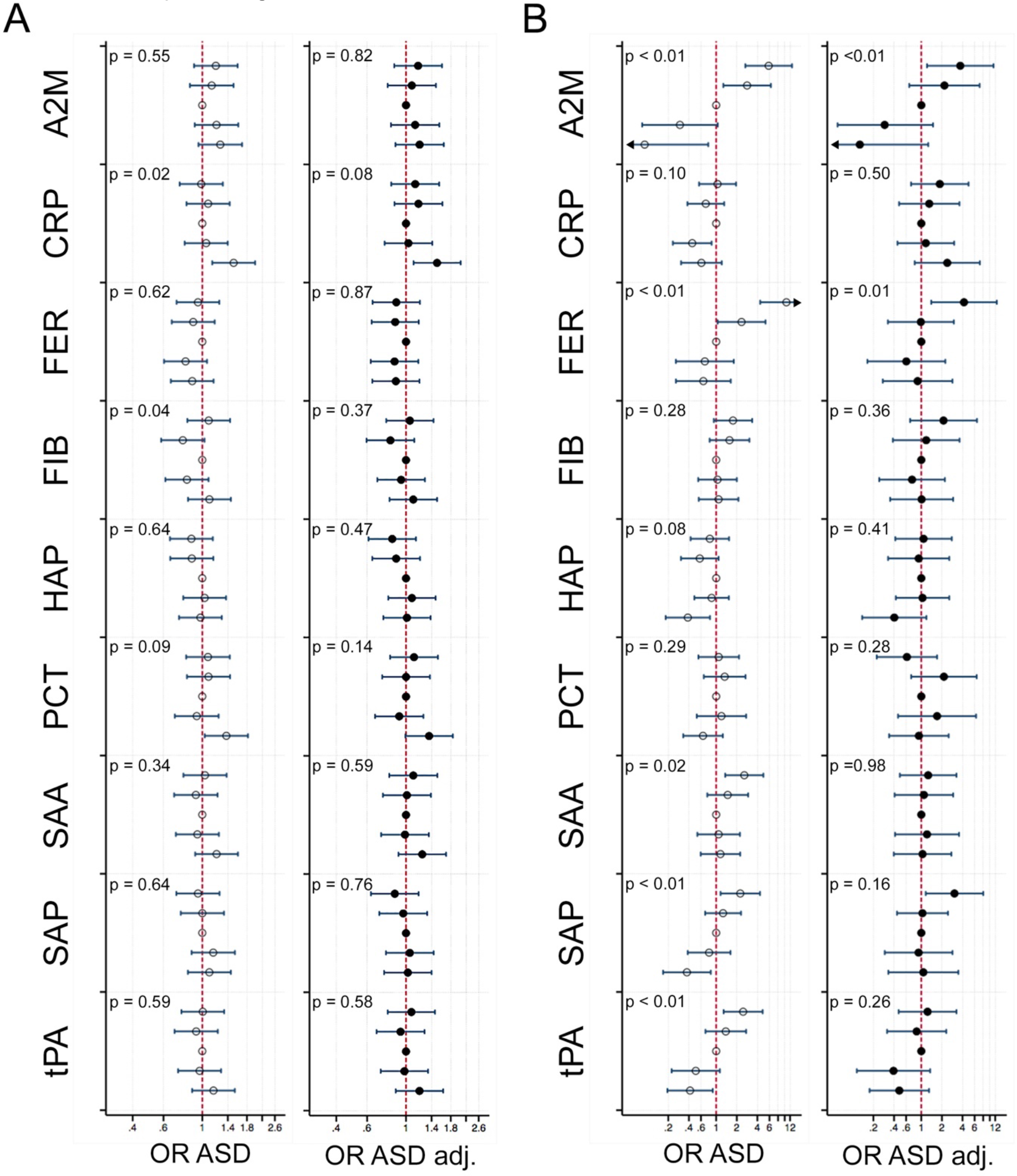
The relationship between APP and odds of ASD when comparing 924 ASD cases to 1092 unaffected individuals selected from the cohort (**A**) and when comparing 203 ASD cases to their matched, unaffected siblings (**B**). Quintiles of each APP were created using the distribution of z-scores among unaffected individuals to set the cut-offs and the middle quintile was used as the referent category. Results are shown for unadjusted estimates (open circles) and fully adjusted estimates (solid circles). Models were adjusted for maternal age, psychiatric history, country of origin, and hospitalization for infection during pregnancy; children’s birth order, sex, gestational age at birth, size for gestational age, and mode of delivery. P-values are shown for a Wald test with a null hypothesis that all APP categorical terms were jointly equal to zero, as a test of whether each APP was generally associated with the outcome. Truncated confidence intervals in the matched sibling comparison are indicated with an arrow. Abbreviations: **A2M**: α-2 macroglobulin; **CRP**: C reactive protein; **FER**: ferritin; **FIB**: fibrinogen; **HAP**: haptoglobin; **PCT**: procalcitonin; **SAA**: serum amyloid A; **SAP**: serum amyloid P; and **tPA**: tissue plasminogen activator.

**Figure S7.**
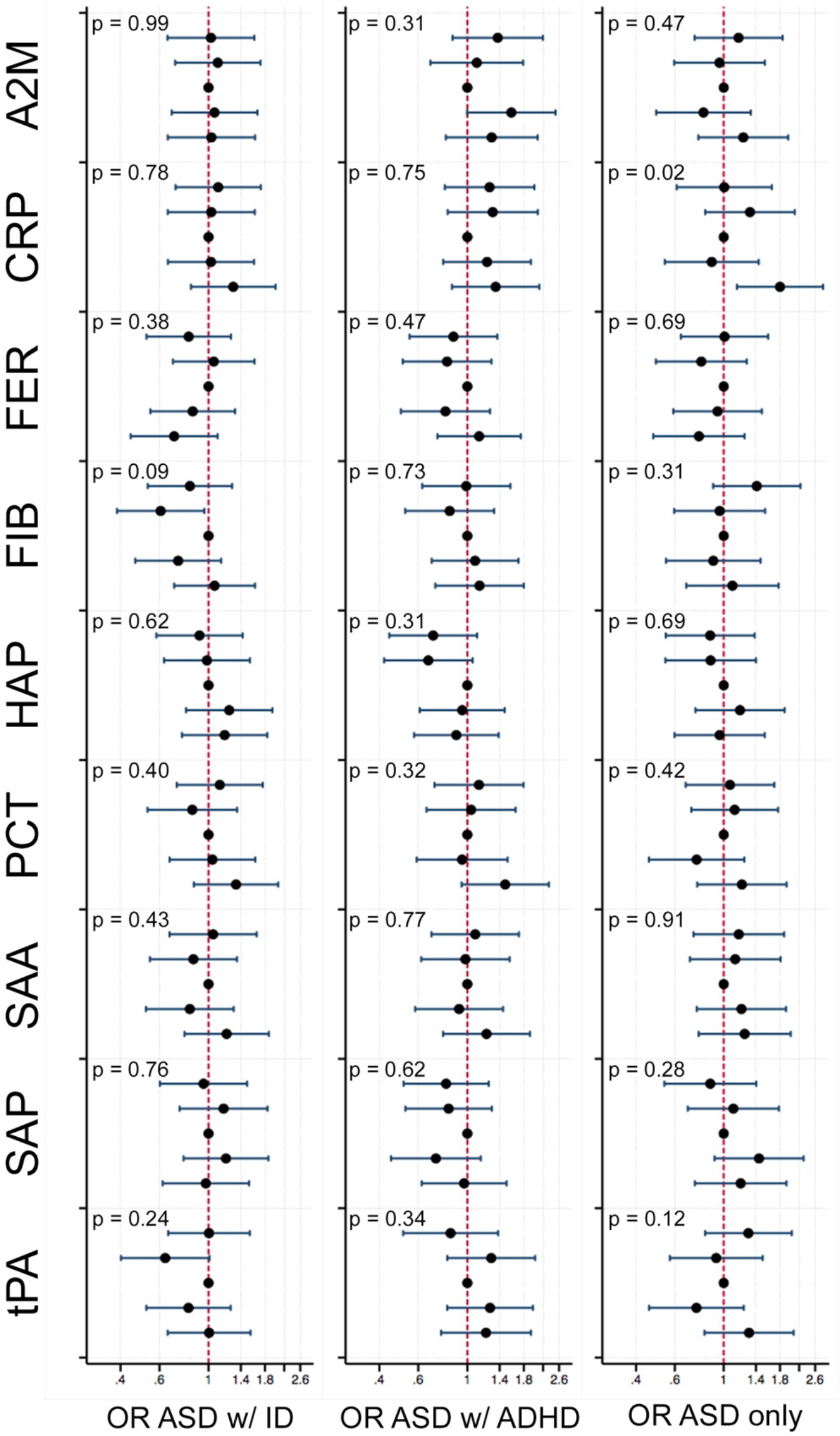
The relationship between APP and odds of ASD when the outcome is stratified by co-occurring ID and ADHD. Results are shown for ASD with co-morbid ID, ASD with co-morbid ADHD, and ASD without co-occurring ID or ADHD (“ASD only”), compared to 1092 unaffected individuals selected from the cohort. Quintiles of each APP were created using the distribution of z-scores among unaffected individuals to set the cut-offs and the middle quintile was used as the referent category. Models were adjusted for maternal age, psychiatric history, country of origin, and hospitalization for infection during pregnancy; children’s birth order, sex, gestational age at birth, size for gestational age, and mode of delivery. P-values are shown for a Wald test with a null hypothesis that all APP categorical terms were jointly equal to zero, as a test of whether each APP was generally associated with the outcome. Abbreviations: **A2M**: α-2 macroglobulin; **CRP**: C reactive protein; **FER**: ferritin; **FIB**: fibrinogen; **HAP**: haptoglobin; **PCT**: procalcitonin; **SAA**: serum amyloid A; **SAP**: serum amyloid P; and **tPA**: tissue plasminogen activator.

**Figure S8.**
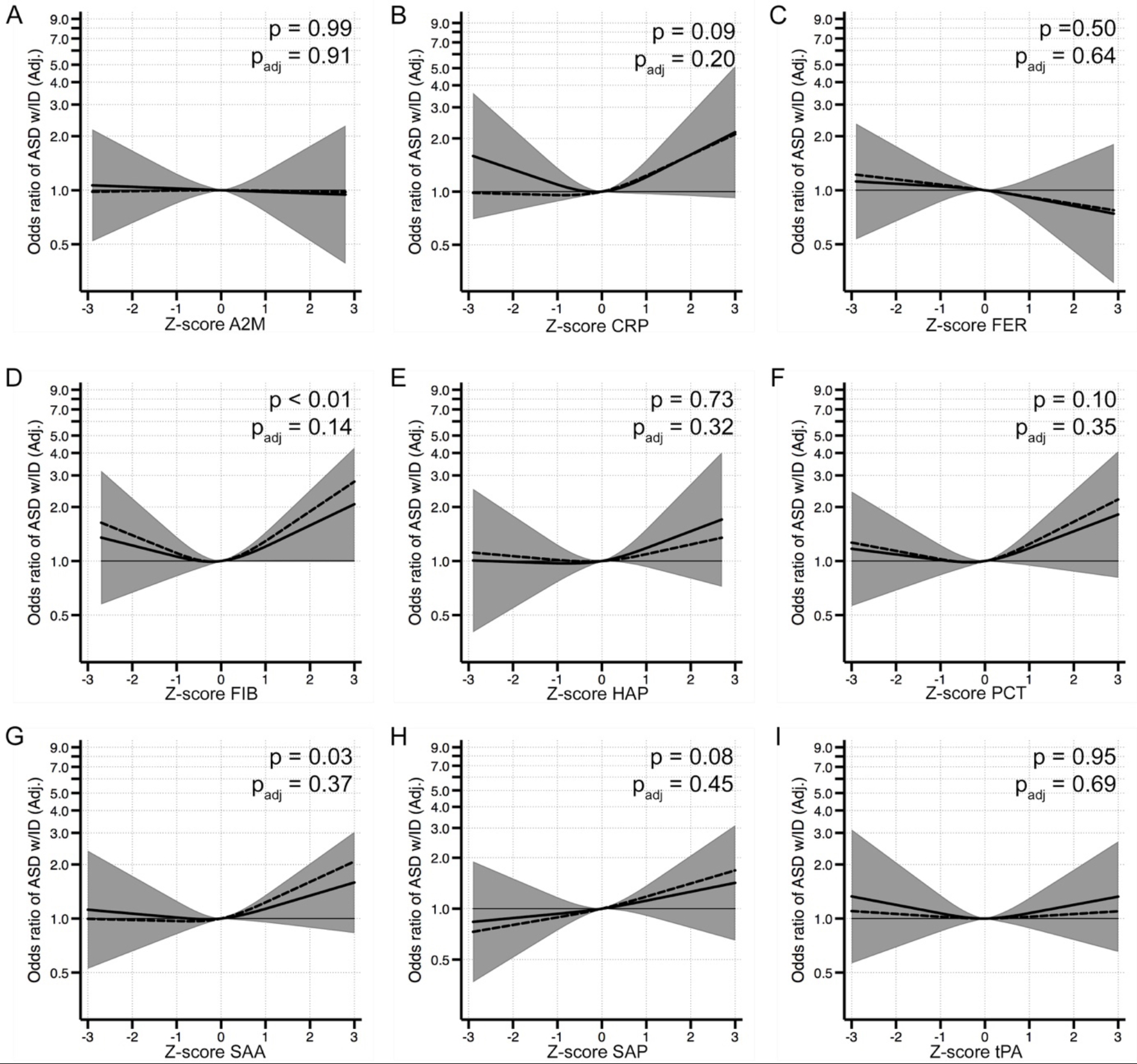
The relationship between APP and odds of ASD with co-occurring ID when comparing 321 individuals affected by ASD with co-occurring ID to 1092 unaffected individuals selected from the cohort. Each panel displays the odds of ASD according to APP z-score, flexibly fit using restricted cubic spline models with three knots and a z-score=0 as the referent. The dashed line represents the unadjusted estimate of the relationship between each APP and odds of ASD. The solid line represents the fully adjusted model, adjusted for maternal age, psychiatric history, country of origin, and hospitalization for infection during pregnancy; children’s birth order, sex, gestational age at birth, size for gestational age, and mode of delivery. The gray bands represent the 95% confidence interval for the fully adjusted model. P-values are shown for a Wald test with a null hypothesis that all APP spline terms were jointly equal to zero, as a test of whether each APP was generally associated with the outcome. P values are shown for spline terms in both the unadjusted models (p) and the adjusted models (p_adj_). Abbreviations: **A2M**: α-2 macroglobulin; **CRP**: C reactive protein; **FER**: ferritin; **FIB**: fibrinogen; **HAP**: haptoglobin; **PCT**: procalcitonin; **SAA**: serum amyloid A; **SAP**: serum amyloid P; and **tPA**: tissue plasminogen activator.

**Figure S9.**
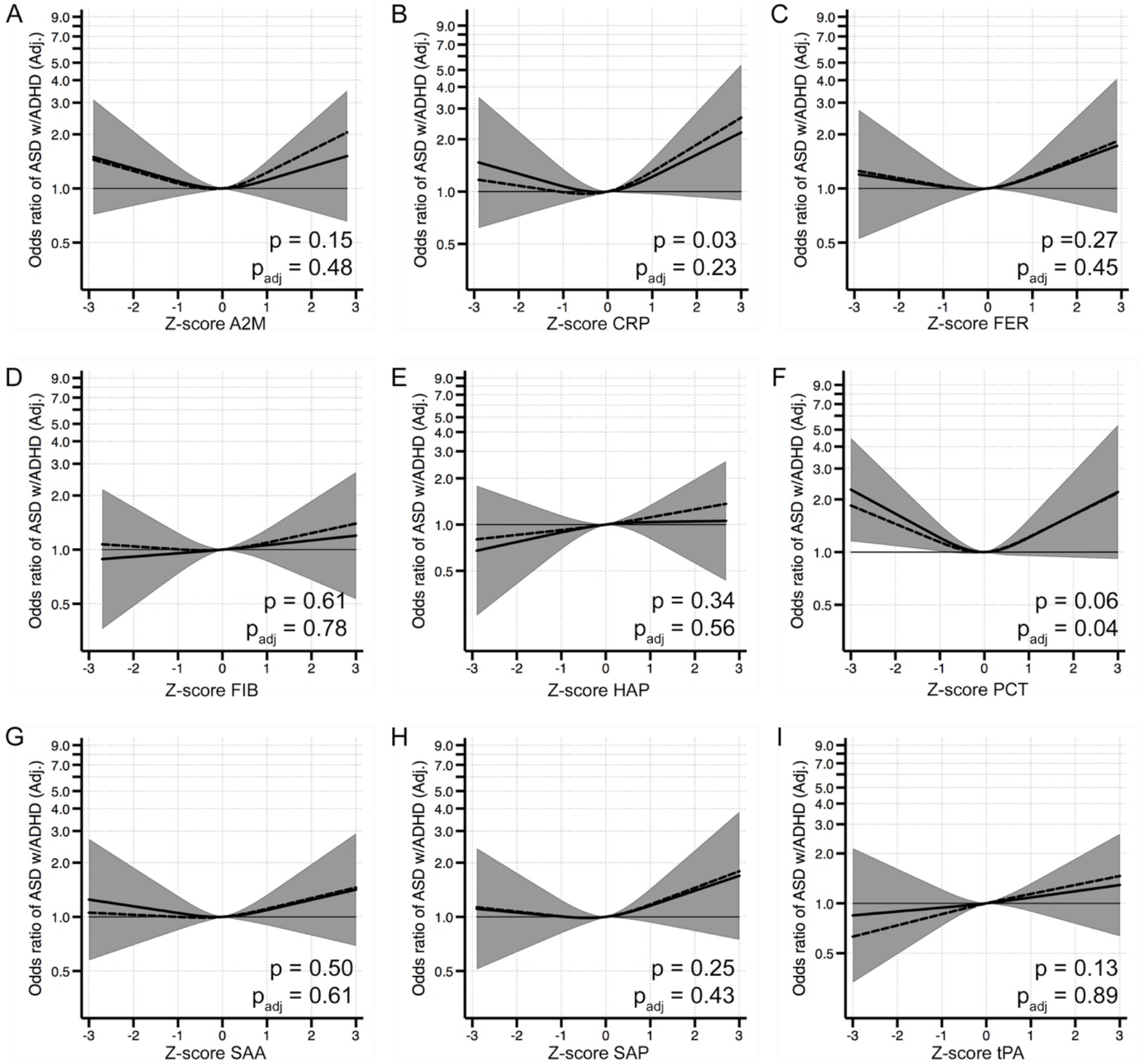
The relationship between APP and odds of ASD with co-occurring ADHD when comparing 322 individuals affected by ASD with co-occurring ADHD to 1092 unaffected individuals selected from the cohort. Each panel displays the odds of ASD according to APP z-score, flexibly fit using restricted cubic spline models with three knots and z-score=0 as the referent. The dashed line represents the unadjusted estimate of the relationship between each APP and odds of ASD. The solid line represents the fully adjusted model, adjusted for maternal age, psychiatric history, country of origin, and hospitalization for infection during pregnancy; children’s birth order, sex, gestational age at birth, size for gestational age, and mode of delivery. The gray bands represent the 95% confidence interval for the fully adjusted model. P-values are shown for a Wald test with a null hypothesis that all APP spline terms were jointly equal to zero, as a test of whether each APP was generally associated with the outcome. P values are shown for spline terms in both the unadjusted models (p) and the adjusted models (p_adj_). Abbreviations: **A2M**: α-2 macroglobulin; **CRP**: C reactive protein; **FER**: ferritin; **FIB**: fibrinogen; **HAP**: haptoglobin; **PCT**: procalcitonin; **SAA**: serum amyloid A; **SAP**: serum amyloid P; and **tPA**: tissue plasminogen activator.

**Figure S10.**
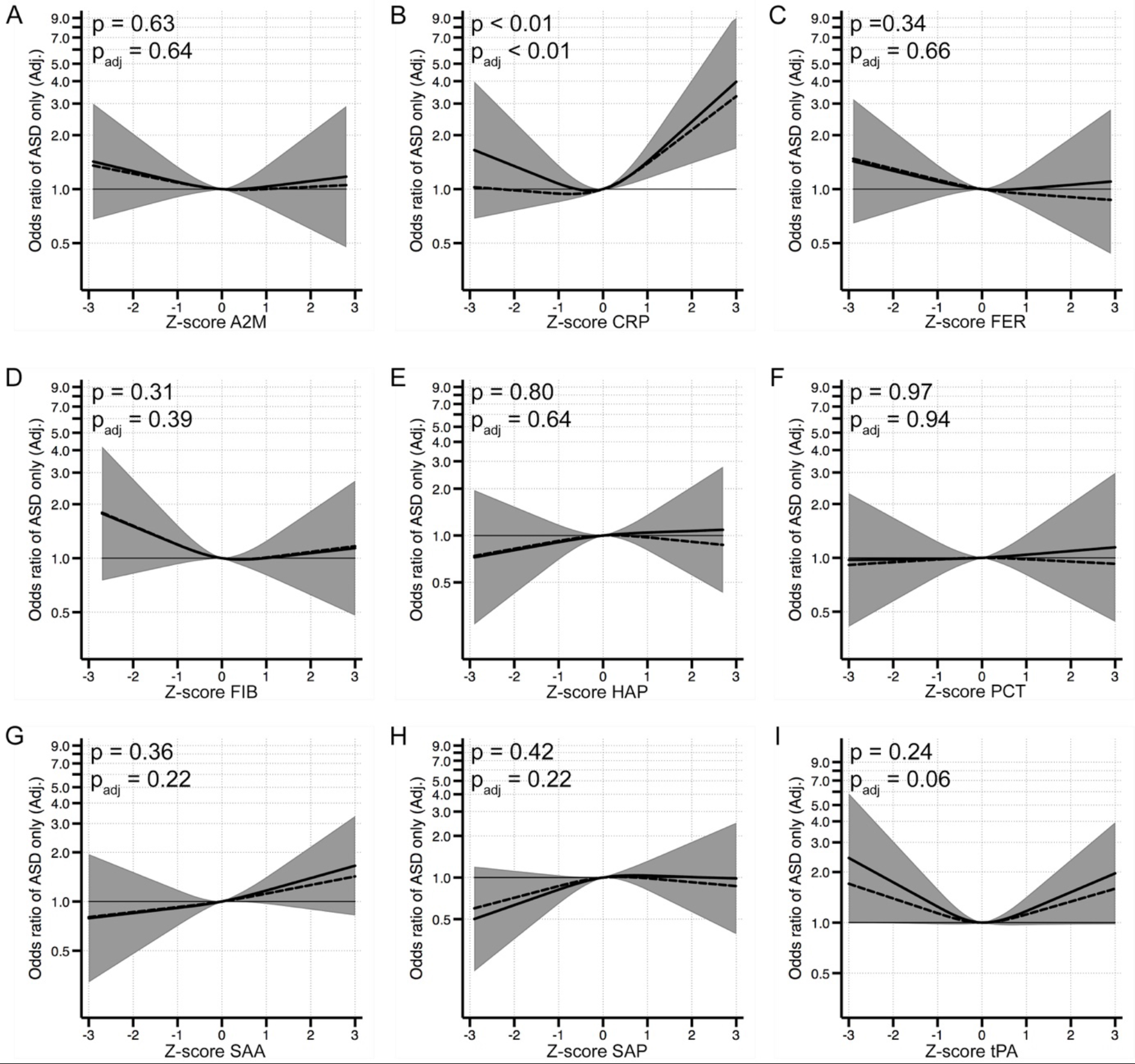
The relationship between APP and odds of ASD without co-occurring ID or ADHD (“ASD only”) when comparing 281 individuals affected by ASD and without a diagnosis of co-occurring ID or ADHD to 1092 unaffected individuals selected from the cohort. Each panel displays the odds of ASD according to APP z-score, flexibly fit using restricted cubic spline models with three knots and a z-score=0 as the referent. The dashed line represents the unadjusted estimate of the relationship between each APP and odds of ASD. The solid line represents the fully adjusted model, adjusted for maternal age, psychiatric history, country of origin, and hospitalization for infection during pregnancy; children’s birth order, sex, gestational age at birth, size for gestational age, and mode of delivery. The gray bands represent the 95% confidence interval for the fully adjusted model. P-values are shown for a Wald test with a null hypothesis that all APP spline terms were jointly equal to zero, as a test of whether each APP was generally associated with the outcome. P values are shown for spline terms in both the unadjusted models (p) and the adjusted models (p_adj_). Abbreviations: **A2M**: α-2 macroglobulin; **CRP**: C reactive protein; **FER**: ferritin; **FIB**: fibrinogen; **HAP**: haptoglobin; **PCT**: procalcitonin; **SAA**: serum amyloid A; **SAP**: serum amyloid P; and **tPA**: tissue plasminogen activator.

**Figure S11.**
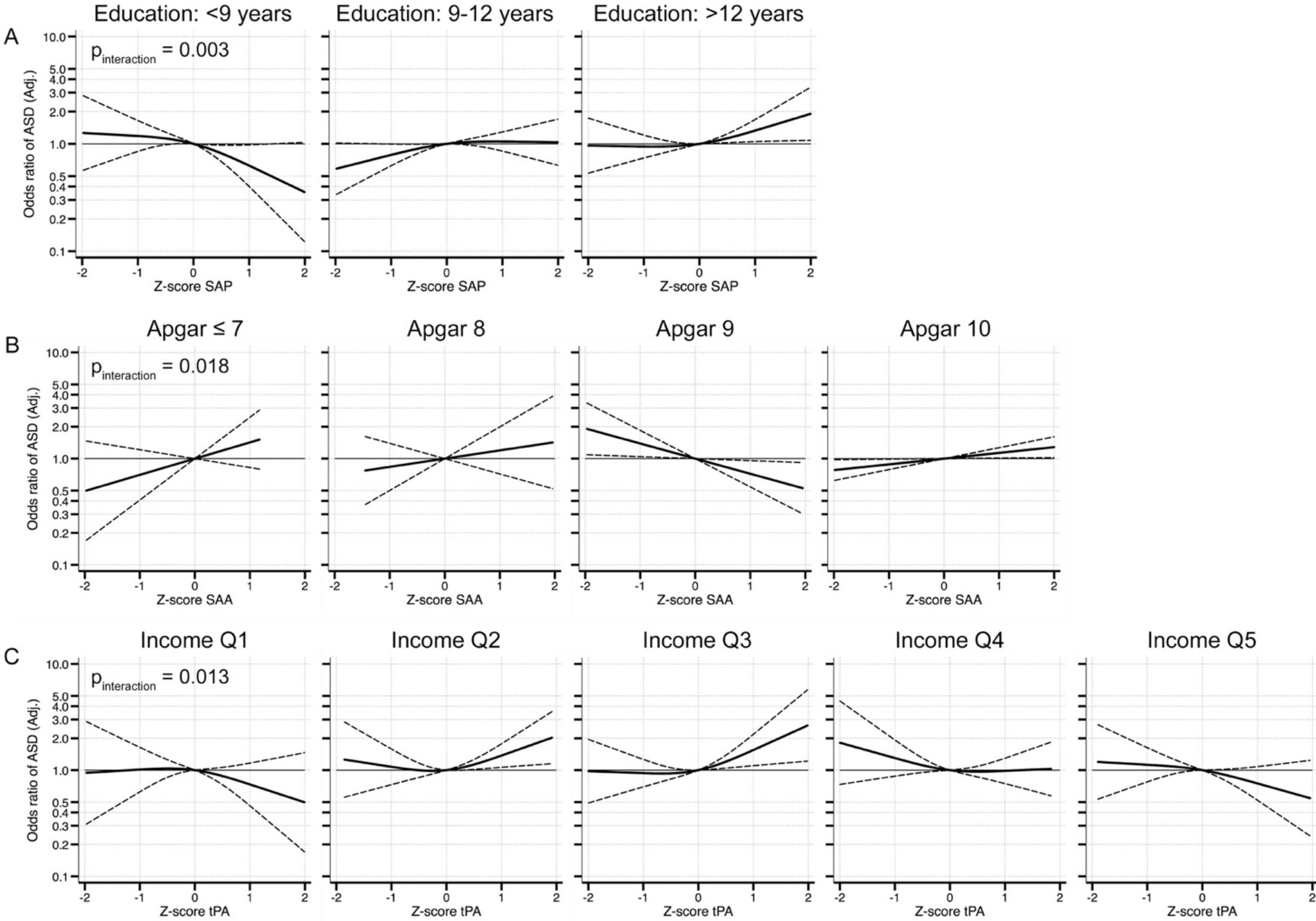
Interaction models for categorical variables. Odds ratios for ASD over the range of APP are shown separately according to strata of parental education level **(A)**, Apgar score at 5 minutes **(B)**, and parental income at birth **(C)**. Solid lines represent the OR estimate for each group and dashed lines represent the 95% confidence interval for the fully adjusted model. P-values for interaction (a likelihood ratio test comparing a model with interaction terms to a model without interaction terms) are shown. Abbreviations: **SAA**: serum amyloid A; **SAP**: serum amyloid P; and **tPA**: tissue plasminogen activator.

**Figure S12.**
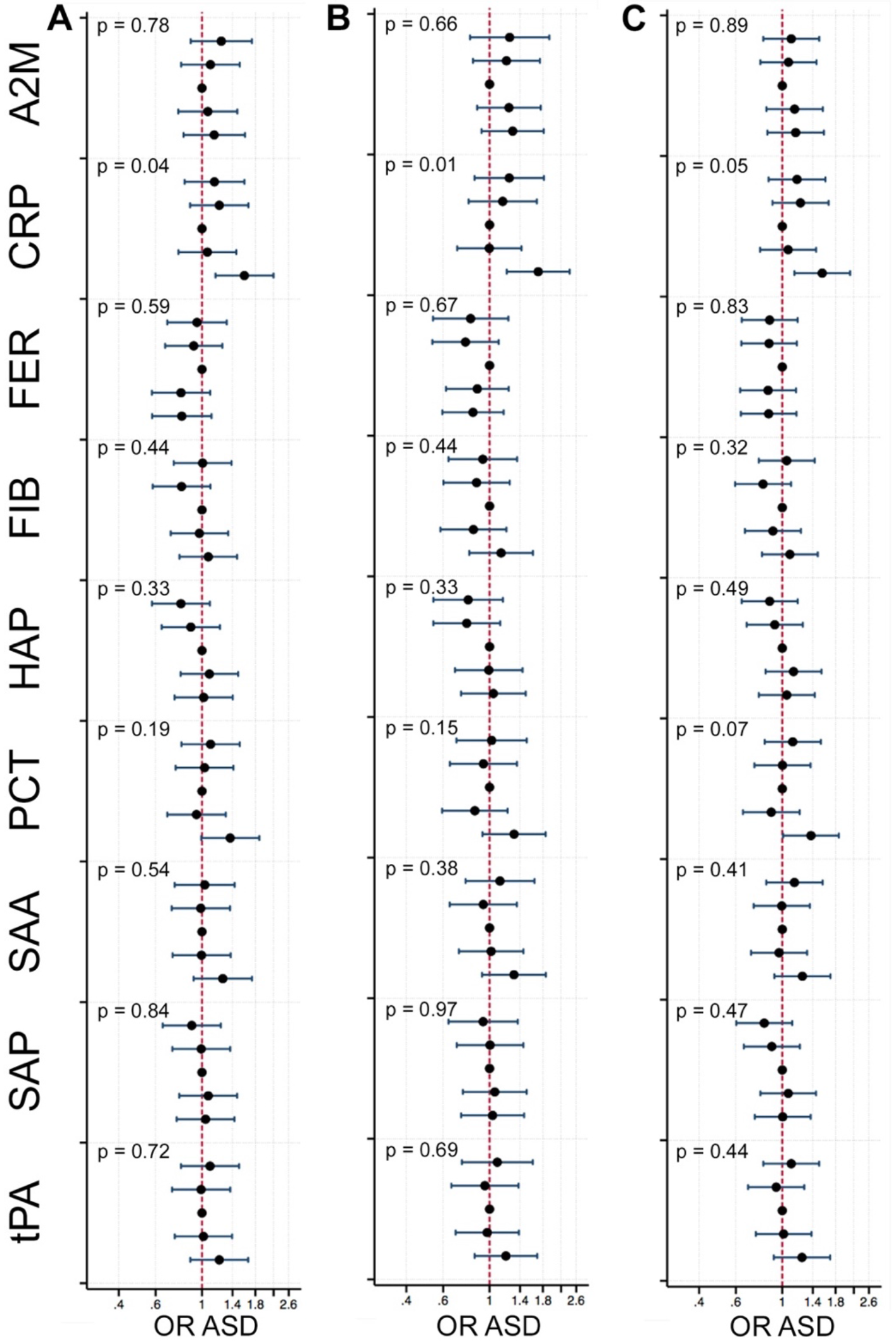
The relationship between APP and odds of ASD in sensitivity analyses. In (**A**), 924 ASD cases are compared to 1092 unaffected individuals selected from the cohort, adjusting for birth year, total protein content of the eluate from the neonatal dried blood spot sample, and age (in days) at neonatal blood sample, in addition to the covariates of the fully adjusted model (maternal age, psychiatric history, country of origin, and hospitalization for infection during pregnancy; children’s birth order, sex, gestational age at birth, size for gestational age, and mode of delivery). In (**B**), individuals with a total protein concentration in the lowest quartile were excluded, and results are shown for the fully adjusted model comparing 688 ASD cases to 819 unaffected individuals selected from the cohort. In (**C**), results are shown for the fully adjusted model using a multi-level modeling approach that accounts for the clustering of APP observations in elution batches and multiplex analysis plate, comparing 924 ASD cases to 1092 unaffected individuals. We used generalized linear and mixed models to fit a 3-level random intercept model with logit link, allowing for the intercepts of both plates and batch to vary, but assuming that the relationship between the log-odds of ASD and each APP does not vary across plates or batches.

### Supplemental Tables

**Table S1.** Diagnostic codes used in the ascertainment of ASD, ADHD, and ID in the Stockholm Youth Cohort.

| **Coding system** | **Autism Spectrum Disorders (ASD)** | **Attention Deficit/Hyperactivity Disorder (ADHD)** | **Intellectual Disability (ID)** |
| --- | --- | --- | --- |
| ICD-10<sup>a,b,c</sup> | F84 | F90 | F70-F79 |
| DSM-IV<sup>b</sup> | 299 | 314 | 317-319 |
| Other | Habilitation Register<sup>d</sup> | Prescription Drug Register<sup>e</sup>:<br>methylphenidate [N06BA04] or atomoxetine [N06BA09] | Habilitation Register<sup>d</sup> |

<sup>a</sup> The National Patient Register (NPR): including inpatient care from 1973, outpatient physician visits in specialist care from 1997, and children and adolescent psychiatry from 2006.
<sup>b</sup> Stockholm Clinical Database for Child and Adolescent Psychiatry: DSM-IV was used until 2008; ICD-10 was used thereafter.
<sup>c</sup> VAL register: Stockholm county health care databases including in- and outpatient care, regardless of specialty/primary health care.
<sup>d</sup> The Habilitation Register (HAB): Provides data on utilization of Stockholm County Habilitation services according to type of disability, such as pervasive developmental disorders and intellectual disability (recorded as present or absent). HAB register was used until 2008; since 2008 it has been included in the VAL databases as ICD-coded diagnoses (see C above).
<sup>e</sup> The Prescription Drug Register (PDR) contains data on medications dispensed to the entire population in Sweden since 1 July 2005. Receipt of a prescription for ADHD medications is a useful proxy for an ADHD diagnosis, as Swedish medical guidelines mandate that ADHD medications should only be prescribed by a psychiatric specialist and after other (non-pharmacological) interventions have failed.

**Table S2.** Quality control statistics for multiplex assays to analyze acute phase protein concentrations in neonatal dried blood spots.

| Analyte | %CV (controls)<sup>a</sup> | Below LLOQ | LLOQ<sup>b</sup> | Above ULOQ | ULOQ<sup>c</sup> | Imputed<sup>d</sup> |
| --- | --- | --- | --- | --- | --- | --- |
| α-2-macroglobulin | 11.5 | 0 (0.00%) | 0.40 ng/ml | 0 (0.00%) | 1658.00 ng/ml | 2 (0.10%) |
| Haptoglobin | 23.3 | 3 (0.15%) | 0.09 ng/ml | 2 (0.10%) | 99.05 ng/ml | 11 (0.55%) |
| C Reactive Protein | 15.4 | 5 (0.25%) | 0.01 ng/ml | 0 (0.00%) | 22.95 ng/ml | 19 (0.94%) |
| Serum Amyloid P | 9.1 | 1 (0.05%) | 0.01 ng/ml | 0 (0.00%) | 185.20 ng/ml | 1 (0.05%) |
| Procalcitonin | 22.3 | 9 (0.45%) | 4.71 pg/ml | 0 (0.00%) | 5430 pg/ml | 95 (4.71%) |
| Ferritin | 11.7 | 0 (0.00%) | 2.39 pg/ml | 0 (0.00%) | 39698.50 pg/ml | 0 (0.00%) |
| Tissue Plasminogen Activator | 23.3 | 4 (0.20%) | 1.07 pg/ml | 0 (0.00%) | 4884 pg/ml | 41 (2.03%) |
| Fibrinogen | 17.0 | 0 (0.00%) | 3.04 ng/ml | 0 (0.00%) | 2911.50 ng/ml | 1 (0.05%) |
| Serum Amyloid A | 22.0 | 0 (0.00%) | 0.50 ng/ml | 0 (0.00%) | 374 ng/ml | 82 (4.07%) |

- The percent CV is based on standardized controls that were run in duplicate on each of the 24 assay plates. - Lower Limit of Quantitation, average across 24 assay plates. - Upper Limit of Quantitation, average across 24 assay plates. - The absolute and relative frequency of observations that were imputed, including those values that were below the LLOQ, above the ULOQ, or that the Bio-plex Manager software indicated were near the LLOQ and thus less certain than values higher in the range of quantitation.

**Table S3.** Characteristics of unaffected and ASD-affected siblings included in the matched sibling comparison.

|  | Unaffected siblings<br>(n = 203) | ASD affected siblings<br>(n = 203) |
| --- | --- | --- |
| **Sex** |  |  |
| Female | 116 (57.1%) | 43 (21.2%) |
| Male | 87 (42.9%) | 160 (78.8%) |
| **Birth Order** |  |  |
| 1st born | 100 (49.3%) | 39 (19.2%) |
| 2nd born | 68 (33.5%) | 105 (51.7%) |
| 3rd or higher | 29 (14.3%) | 41 (20.2%) |
| Missing | 6 (3.0%) | 18 (8.9%) |
| **Mode of Delivery** |  |  |
| Unassisted VD | 129 (63.5%) | 127 (62.6%) |
| Induced VD | 10 (4.9%) | 13 (6.4%) |
| Assisted VD | 19 (9.4%) | 12 (5.9%) |
| Elective CS | 15 (7.4%) | 13 (6.4%) |
| Emergency CS | 24 (11.8%) | 20 (9.9%) |
| Missing | 6 (3.0%) | 18 (8.9%) |
| **Gestational Age** |  |  |
| ≤36 weeks | 7 (3.4%) | 8 (3.9%) |
| 37-41 weeks | 173 (85.2%) | 165 (81.3%) |
| ≥42 weeks | 17 (8.4%) | 12 (5.9%) |
| Missing | 6 (3.0%) | 18 (8.9%) |
| **Size for Gestational Age** |  |  |
| Normal size for GA | 182 (89.7%) | 172 (84.7%) |
| Small for GA | 9 (4.4%) | 4 (2.0%) |
| Large for GA | 6 (3.0%) | 7 (3.4%) |
| Missing | 6 (3.0%) | 20 (9.9%) |
| **Apgar Score at 5 Minutes** |  |  |
| 10 | 158 (77.8%) | 152 (74.9%) |
| 9 | 24 (11.8%) | 24 (11.8%) |
| 8 | 9 (4.4%) | 4 (2.0%) |
| ≤7 | 5 (2.5%) | 4 (2.0%) |
| Missing | 7 (3.4%) | 19 (9.4%) |
| **Age at Neonatal Blood Sample (Days)** |  |  |
| ≤3 | 79 (38.9%) | 66 (32.5%) |
| 4 | 73 (36.0%) | 73 (36.0%) |
| 5 | 16 (7.9%) | 43 (21.2%) |
| 6 | 20 (9.9%) | 11 (5.4%) |
| ≥7 | 13 (6.4%) | 10 (4.9%) |
| Missing | 2 (1.0%) | 0 (0.0%) |
| **Maternal Age (Years)** |  |  |
| >25 | 35 (17.2%) | 25 (12.3%) |
| 25-29 | 67 (33.0%) | 71 (35.0%) |
| 30-34 | 75 (36.9%) | 64 (31.5%) |
| 35-39 | 20 (9.9%) | 38 (18.7%) |
| ≥40 | 6 (3.0%) | 5 (2.5%) |
| **Maternal Psychiatric History** |  |  |
| No | 115 (56.7%) | 114 (56.2%) |
| Yes | 88 (43.3%) | 89 (43.8%) |
| **Maternal BMI** |  |  |
| Normal | 94 (46.3%) | 80 (39.4%) |
| Underweight | 3 (1.5%) | 2 (1.0%) |
| Overweight | 41 (20.2%) | 35 (17.2%) |
| Obese | 15 (7.4%) | 13 (6.4%) |
| Missing | 50 (24.6%) | 73 (36.0%) |
| **Mother Born Outside Sweden** |  |  |
| No | 130 (64.0%) | 130 (64.0%) |
| Yes | 73 (36.0%) | 73 (36.0%) |
| **Family Income Quintile** |  |  |
| 1<sup>st</sup> (Lowest) | 28 (13.8%) | 28 (13.8%) |
| 2<sup>nd</sup> | 47 (23.2%) | 50 (24.6%) |
| **3<sup>rd</sup>** | 45 (22.2%) | 56 (27.6%) |
| **4<sup>th</sup>** | 44 (21.7%) | 45 (22.2%) |
| **5<sup>th</sup>** | 38 (18.7%) | 24 (11.8%) |
| **Parental Education** |  |  |
| **<9 years** | 41 (20.2%) | 41 (20.2%) |
| **9-12 years** | 92 (45.3%) | 92 (45.3%) |
| **>12 years** | 70 (34.5%) | 70 (34.5%) |
| **Maternal Hospitalization for Infection (3rd trimester)** |  |  |
| **No** | 190 (93.6%) | 176 (86.7%) |
| **Yes** | 7 (3.4%) | 9 (4.4%) |
| **Missing** | 6 (3.0%) | 18 (8.9%) |
| **Maternal Anemia During Pregnancy** |  |  |
| **No** | 191 (94.1%) | 192 (94.6%) |
| **Yes** | 12 (5.9%) | 11 (5.4%) |
| **Maternal Dietary Supplementation** |  |  |
| **None** | 95 (46.8%) | 94 (46.3%) |
| **Folic Acid only** | 4 (2.0%) | 1 (0.5%) |
| **Iron only** | 62 (30.5%) | 56 (27.6%) |
| **Multivitamin** | 42 (20.7%) | 52 (25.6%) |
| **Proteins measured in NDBS (median [IQR])** |  |  |
| **A2M (ng/ml)** | 212.4 (34.1, 290.0) | 135.0 (25.0, 223.0) |
| **CRP (ng/ml)** | 1.2 (0.4, 3.8) | 1.2 (0.3, 3.7) |
| **FER (ng/ml)** | 4.8 (3.3, 6.5) | 2.9 (1.9, 4.2) |
| **FIB (ng/ml)** | 282.1 (142.5, 461.9) | 220.4 (117.0, 420.2) |
| **HAP (ng/ml)** | 13.5 (5.1, 78.9) | 18.1 (5.5, 68.7) |
| **PCT (pg/ml)** | 11.2 (8.6, 16.0) | 11.2 (8.6, 15.8) |
| **SAA (ng/ml)** | 2.0 (1.2, 3.9) | 1.7 (1.1, 3.4) |
| **SAP (ng/ml)** | 17.5 (3.2, 37.8) | 13.5 (3.0, 26.9) |
| **tPA (pg/ml)** | 10.6 (8.1, 15.9) | 9.2 (6.5, 13.7) |
| **Total protein (mg/ml)** | 3.4 (3.0, 4.0) | 2.9 (2.4, 3.3) |

Abbreviations: **ASD**: autism spectrum disorders; **ADHD**: attention-deficit/hyperactivity disorder; **ID**: intellectual disability; **VD**: vaginal delivery; **CS**: Caesarean section delivery; **GA**: gestational age; **BMI**: body mass index; **IQR**: interquartile range; **A2M**: α-2 macroglobulin; **CRP**: C reactive protein; **FER**: ferritin; **FIB**: fibrinogen; **HAP**: haptoglobin; **PCT**: procalcitonin; **SAA**: serum amyloid A; **SAP**: serum amyloid P; and **tPA**: tissue plasminogen activator.

**Table S4.** P-values for the association of APP with other covariates among 1092 unaffected individuals. We examined the association of each covariate with the APP z-scores by regressing each APP over the categories of the covariate, followed by a joint Wald test of the hypothesis that all coefficients for the categorical indicators are equal to zero, a test of whether each APP was generally associated with the covariate.

|  | A2M | CRP | FER | FIB | HAP | PCT | SAA | SAP | tPA |
| --- | --- | --- | --- | --- | --- | --- | --- | --- | --- |
| Sex | 0.413 | **0.001** | 0.226 | 0.397 | 0.745 | 0.301 | 0.597 | 0.629 | 0.244 |
| Birth Order | 0.290 | **0.002** | 0.109 | 0.571 | 0.359 | 0.051 | 0.184 | **0.030** | **0.012** |
| Birth Year | 0.179 | 0.754 | **0.001** | 0.077 | 0.956 | 0.420 | 0.835 | 0.338 | 0.055 |
| Mode | 0.071 | **<0.001** | **0.040** | 0.844 | **<0.001** | 0.521 | **0.039** | **0.039** | 0.959 |
| Gestational Age | **0.043** | **<0.001** | 0.088 | 0.265 | **0.013** | 0.414 | 0.527 | **0.000** | 0.298 |
| Size for GA | 0.458 | 0.350 | 0.425 | 0.926 | **0.006** | 0.688 | 0.243 | 0.756 | 0.796 |
| Apgar | 0.266 | 0.170 | 0.677 | 0.202 | 0.466 | **0.023** | 0.258 | 0.303 | 0.933 |
| Sample Age | 0.359 | **<0.001** | 0.515 | **0.011** | **<0.001** | **<0.001** | **<0.001** | 0.294 | **0.017** |
| Maternal Age | 0.447 | 0.174 | 0.166 | 0.118 | 0.175 | 0.275 | 0.672 | 0.769 | **0.006** |
| Maternal Psych. Hist. | 0.919 | 0.067 | 0.371 | 0.189 | 0.798 | 0.887 | 0.509 | 0.265 | 0.951 |
| Maternal BMI | 0.499 | 0.572 | 0.777 | 0.164 | 0.937 | 0.201 | 0.251 | 0.059 | 0.448 |
| Maternal Migration | 0.552 | 0.620 | 0.300 | **0.037** | 0.353 | 0.173 | **0.030** | **0.003** | 0.266 |
| Income | 0.802 | 0.156 | 0.739 | 0.082 | 0.083 | **0.020** | **<0.001** | 0.785 | **0.012** |
| Education | 0.842 | 0.298 | 0.777 | 0.584 | 0.796 | 0.388 | 0.883 | 0.777 | 0.681 |
| Maternal Infection | 0.172 | 0.349 | 0.955 | **0.006** | 0.683 | 0.052 | 0.174 | 0.569 | **0.010** |
| Anemia | 0.403 | 0.658 | 0.358 | 0.078 | 0.926 | **0.025** | 0.672 | 0.256 | 0.322 |
| Supplement | 0.207 | 0.476 | 0.360 | 0.054 | 0.199 | 0.092 | **0.022** | 0.117 | **0.045** |

Abbreviations: **GA**: gestational age; **Psych**: Psychiatric; **BMI**: body mass index; **A2M**: α-2 macroglobulin; **CRP**: C reactive protein; **FER**: ferritin; **FIB**: fibrinogen; **HAP**: haptoglobin; **PCT**: procalcitonin; **SAA**: serum amyloid A; **SAP**: serum amyloid P; and **tPA**: tissue plasminogen activator.

**Table S5.**
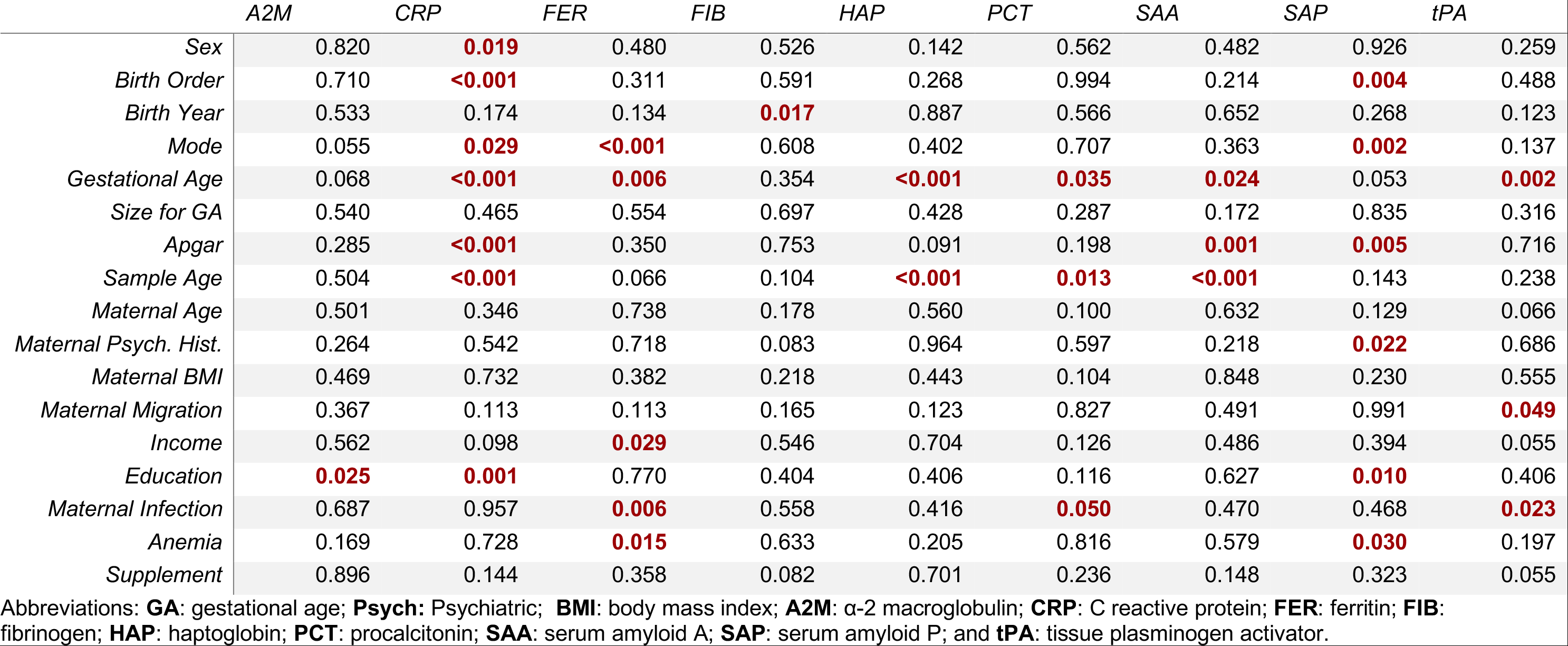
P-values for the association of APP with other covariates among 924 ASD affected individuals. We examined the association of each covariate with the APP z-scores by regressing each APP over the categories of the covariate, followed by a joint Wald test of the hypothesis that all coefficients for the categorical indicators are equal to zero, a test of whether each APP was generally associated with the covariate.

**Table S6.**
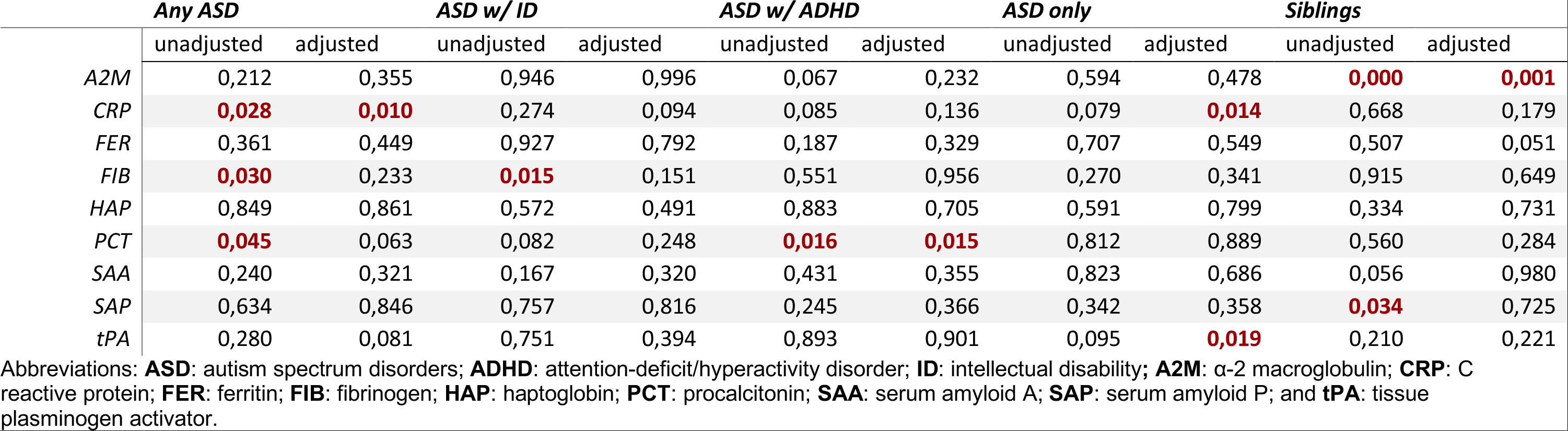
P-values for non-linearity in restricted cubic spline analyses. Not all relationships we considered are necessarily non-linear. To assess evidence for non-linear relationships, we tested the null-hypothesis that all spline terms that would indicate a change in the slope of the relationship (i.e., all but the first spline term) are equal to zero using a Wald test.

|  | *Any ASD* |  | *ASD w/ ID* |  | *ASD w/ ADHD* |  | *ASD only* |  | *Siblings* |  |
| --- | --- | --- | --- | --- | --- | --- | --- | --- | --- | --- |
|  | unadjusted | adjusted | unadjusted | adjusted | unadjusted | adjusted | unadjusted | adjusted | unadjusted | adjusted |
| *A2M* | 0,212 | 0,355 | 0,946 | 0,996 | 0,067 | 0,232 | 0,594 | 0,478 | **0,000** | **0,001** |
| *CRP* | **0,028** | **0,010** | 0,274 | 0,094 | 0,085 | 0,136 | 0,079 | **0,014** | 0,668 | 0,179 |
| *FER* | 0,361 | 0,449 | 0,927 | 0,792 | 0,187 | 0,329 | 0,707 | 0,549 | 0,507 | 0,051 |
| *FIB* | **0,030** | 0,233 | **0,015** | 0,151 | 0,551 | 0,956 | 0,270 | 0,341 | 0,915 | 0,649 |
| *HAP* | 0,849 | 0,861 | 0,572 | 0,491 | 0,883 | 0,705 | 0,591 | 0,799 | 0,334 | 0,731 |
| *PCT* | **0,045** | 0,063 | 0,082 | 0,248 | **0,016** | **0,015** | 0,812 | 0,889 | 0,560 | 0,284 |
| *SAA* | 0,240 | 0,321 | 0,167 | 0,320 | 0,431 | 0,355 | 0,823 | 0,686 | 0,056 | 0,980 |
| *SAP* | 0,634 | 0,846 | 0,757 | 0,816 | 0,245 | 0,366 | 0,342 | 0,358 | **0,034** | 0,725 |
| *tPA* | 0,280 | 0,081 | 0,751 | 0,394 | 0,893 | 0,901 | 0,095 | **0,019** | 0,210 | 0,221 |

Abbreviations: **ASD**: autism spectrum disorders; **ADHD**: attention-deficit/hyperactivity disorder; **ID**: intellectual disability; **A2M**: α-2 macroglobulin; **CRP**: C reactive protein; **FER**: ferritin; **FIB**: fibrinogen; **HAP**: haptoglobin; **PCT**: procalcitonin; **SAA**: serum amyloid A; **SAP**: serum amyloid P; and **tPA**: tissue plasminogen activator.

**Table S7.** P-values for interaction using a likelihood ratio test comparing a model without interaction terms nested in model with interaction terms. A low p-value on this test indicates evidence to reject the null hypothesis of the absence of interaction between the APP and the covariates tested.

|  | **A2M** | **CRP** | **FER** | **FIB** | **HAP** | **PCT** | **SAA** | **SAP** | **tPA** |
| --- | --- | --- | --- | --- | --- | --- | --- | --- | --- |
| Sex | 0.473 | 0.726 | 0.097 | 0.746 | 0.360 | 0.340 | 0.975 | 0.385 | 0.239 |
| Birth order | 0.869 | 0.296 | 0.103 | 0.281 | 0.764 | 0.345 | 0.856 | 0.179 | 0.371 |
| Mode | 0.887 | 0.474 | 0.240 | 0.991 | 0.202 | 0.801 | 0.226 | 0.520 | 0.640 |
| Gestational age | 0.929 | 0.556 | 0.860 | 0.583 | 0.872 | 0.455 | 0.153 | 0.165 | 0.125 |
| Size for gestational age | 0.188 | 0.607 | 0.810 | 0.621 | 0.341 | 0.640 | 0.600 | 0.365 | 0.223 |
| Apgar score | 0.312 | 0.241 | 0.784 | 0.543 | 0.233 | 0.867 | **0.018** | 0.674 | 0.415 |
| Maternal Age | 0.872 | 0.593 | 0.779 | 0.473 | 0.449 | 0.718 | 0.999 | **0.026** | 0.352 |
| Maternal Psychiatric History | 0.331 | **0.037** | 0.619 | 0.093 | 0.710 | 0.911 | 0.369 | 0.293 | 0.575 |
| Maternal BMI | 0.810 | 0.112 | 0.408 | 0.298 | 0.828 | 0.710 | 0.801 | 0.485 | 0.989 |
| Maternal Migration | 0.538 | 0.235 | 0.612 | 0.440 | 0.352 | 0.321 | 0.131 | 0.143 | 0.778 |
| Parental Income | 0.515 | 0.410 | 0.098 | 0.640 | 0.956 | 0.242 | 0.157 | 0.387 | **0.013** |
| Parental Education | 0.070 | 0.215 | 0.321 | 0.737 | 0.399 | 0.849 | 0.707 | **0.003** | 0.174 |
| Maternal Infection | **0.008** | 0.511 | **0.043** | **0.017** | 0.799 | **0.016** | 0.218 | 0.741 | **0.001** |
| Maternal Anemia | 0.100 | 0.864 | **0.010** | 0.185 | 0.526 | 0.204 | 0.816 | **0.043** | 0.311 |
| Maternal Supplement Use | 0.907 | 0.283 | 0.631 | 0.500 | 0.546 | 0.116 | 0.078 | 0.565 | 0.917 |

Abbreviations: **A2M**: α-2 macroglobulin; **CRP**: C reactive protein; **FER**: ferritin; **FIB**: fibrinogen; **HAP**: haptoglobin; **PCT**: procalcitonin; **SAA**: serum amyloid A; **SAP**: serum amyloid P; and **tPA**: tissue plasminogen activator

